# PrinTE: A Forward Simulation Framework for Studying the Role of Transposable Elements in Genome Expansion and Contraction

**DOI:** 10.1101/2025.06.04.657780

**Authors:** Christopher W. Benson, Ting-Hsuan Chen, Susan J. Thomson, Cecilia H. Deng, Shujun Ou

## Abstract

Genome expansion and contraction are reportedly driven by transposable element (TE) activity, but the underlying dynamics remain enigmatic due to a lack of historical records tracing these changes. Here, we present PrinTE for versatile, forward-time simulation of whole-genome sequences with highly customizable transposon dynamics. Through simulations, we confirm that the distribution of TE sequence divergence reflects their historical insertion and deletion dynamics, which can be used to infer TE dynamic parameters through PrinTE simulations. We analyzed the pangenome of *Pucciniomycotina*, a subdivision of fungi containing myrtle rust (*Austropuccinia psidii*), which drastically expanded its genome size to 1018 Mb. Our analyses reveal that the best strategy for controlling genome size is to avoid the invasion of LTR retrotransposons (LTR-RTs). While illegitimate recombination (IR) is considered the most effective counteraction of LTR-RT invasions leaving only solo LTR remnants, we observed a strong positive correlation between solo:intact LTR ratio (strength of LTR-RT removal) and genome size (*r* = 0.65), and a near-linear correlation between solo LTR count and genome size (*r* = 0.98). This result suggests that IR alone may not effectively prevent genome obesity. Through simulation of *Pucciniomycotina* genomes, we proposed that *A. psidii* might experience a prolonged period of genome expansion followed by a short, potent, and likely ongoing period of contraction. PrinTE is freely available at https://github.com/cwb14/PrinTE.git.

## Background

Transposable elements (TEs), or “jumping genes”, are dynamic DNA sequences that replicate and relocate within genomes. TEs constitute 46% of the human (*Homo sapiens*) genome ^1^, 85% of the maize (*Zea mays*) genome ^2^, and over 90% of the genome of the fungal pathogen *Austropuccinia psidii* that causes the myrtle rust disease ^3^. The prevalence of TEs across the Tree of Life reflects their dual roles as both genomic parasites and drivers of evolutionary innovation. Beyond inflating genome sizes, TEs drive structural variation, modulate gene expression, and facilitate chromosomal rearrangements ^4^. In both plant and mammalian genomes, bursts of TE activity correlate with speciation events and adaptive radiations, enabling rapid responses to environmental stressors like pathogens or climate shifts ^5^.

Recent studies highlight TE’s contribution to the collective genetic repertoire of a species. Core genomes, conserved across individuals, are often gene-rich and TE-poor ^2^, while “variable” or “dispensable” regions are TE-dense and enriched for stress-response genes, pathogen resistance loci, and structural variants ^6^. In *Brassica oleracea*, resistance gene analogs cluster in TE-rich variable regions ^7^. Similarly, tandem clusters of leucine-rich repeat (LRR) genes in maize are found in genomic regions rich with young long terminal repeat (LTR) retrotransposons (LTR-RT) ^8^, suggesting that TE-driven structural plasticity contributes to adaptive traits. Related lineages can become karyotypically disparate over generations due to differences in their ability to repress TEs ^9^. The global impact of TEs on macro-chromosomal structure plays an important role in evolutionary fates by influencing meiotic stability and similarity between cytotypes ^10^. These findings underscore TEs as dynamic regulators of genome evolution, influencing both microevolutionary adaptation and macroevolutionary diversification.

Genome size varies considerably across taxa. Mammalian and bird genomes appear more tightly constrained in genome size than those of both fungi and plants ^11^. Numerous studies across the Mammalia class indicate this size constraint is not through a lack of TE radiation bursts ^12^ but through mechanistic interplay with purifying selection, where bursts are followed by TE silencing ^13^ and DNA loss ^12^. Genome size variation in plants facilitates adaptation to specific environments, primarily carried out by whole genome duplication (WGD) and TE expansion ^14^, especially LTR-RTs ^15^. Extremes of genome size can be observed from the compact genome of *Arabidopsis thaliana* (∼135 Mb) through to the giant genomes of the Chinese pine (*Pinus tabuliformis*) at ∼25 Gb ^16^, lily (*Lilium sargentiae*) at ∼36 Gb ^17^, and the New Caledonian fork fern (*Tmesipteris oblanceolata*) at ∼160 Gb ^18^. These are examples of unconstrained genome expansion, with extremely high TE content within intergenic regions and introns. Conversely, there are numerous signs that genome size expansion in plants is subject to constraints and regulatory mechanisms. *Arabidopsis* has undergone several WGDs, yet remains compact partially through subsequent gene loss that purges redundant paralogs ^19^. The dynamic of genome size change has been difficult to accurately evaluate primarily because of the lack of high-fidelity tools for simulation and modeling studies.

Advances in long-read sequencing and computational methods have empowered efforts in resolving TE dynamics ^20^, yet reconstructing their evolutionary history remains challenging due to TEs often being fragmented, nested, and highly identical to each other. Simulations of TE evolution are thus indispensable for testing hypotheses about genome diversification, evaluating TE annotation tools, and modeling the interplay between TE activity and selection. By generating synthetic genomes with known TE architectures, including nested insertions, sequence divergence, and polymorphism, researchers can disentangle the stochastic and deterministic forces shaping genome evolution ^21^.

Existing TE simulators vary widely in scope and range of biological scenarios that they mimic. Early models, such as those by Clough et al. (1996), introduced foundational concepts by simulating varying modes of mammalian transposition ^22^. Modern tools address specific niches. SimulaTE ^23^ generates complex TE landscapes through predefined TE architectures, limiting its utility for exploring stochastic evolutionary scenarios. Population-genetic frameworks like SLiM ^24^ excel at modeling selection and demography but compress TEs into individual units, precluding modeling of TE sequence mutagenesis, nested insertion, and the consequential genome size evolution. TESD ^25^ incorporates spatial population structure and selection against deleterious insertions, yet its inaccessibility hinders further application. ReplicaTE ^26^ engineers synthetic genomes by deleting and reinserting TEs into real assemblies but lacks dynamic iterations through evolutionary epochs. Meanwhile, pMEsim ^27^ focuses on simulating polymorphic TE variants for pan-genomes, sacrificing long-term evolutionary dynamics, and simuG ^28^ serves as a generic genome simulator that can simulate TE insertions. GARLIC ^29^ specializes in generating realistic artificial genomic sequences by modeling intergenic sequences based on kmer profiles, GC contents, and local repeat landscape, which serve as high-quality negative controls for evaluating annotation tools. TEgenomeSimulator is a single-timepoint simulator, useful for benchmarking and creating replicas of the TE landscapes of real genomes, but it does not perform forward simulation for complex evolutionary scenarios.

While existing simulation tools collectively address diverse needs, few allow flexibility in genome customization, TE sequence mutagenesis, and dynamic modulation of insertion/deletion rates across evolutionary epochs; none provides granular family-specific controls with iterative simulations for hypothesis testing; and most prioritize synthetic data generation over empirical inference of TE dynamic parameters.

## Results

### Introduction of PrinTE

Here, we present PrinTE, a transposon-focused tool that allows dynamic forward simulation of TE insertions, deletions, truncations, and sequence substitutions. PrinTE enables iterative forward simulations to explore complex genome evolutionary scenarios, including the effects of TE dynamics on genome size evolution, genomic divergence, the impact of selection on TE turnover, and speciation (**Fig. 1**).

**Fig. 1.**
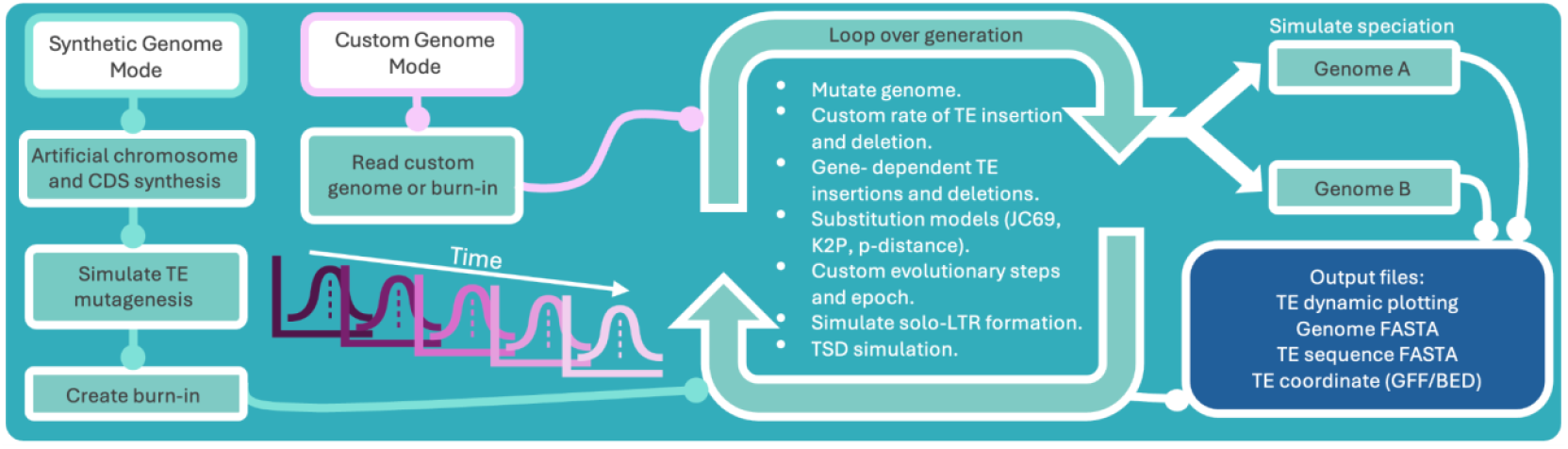
Programmatic components and functionality of PrinTE. The high-level computational workflow of PrinTE is illustrated with rectangles and solid lines. See the main text for detailed descriptions of functionality.

#### Forward simulations of TE insertions, deletions, and genome divergence

PrinTE is a forward-time genome simulation tool designed to model the dynamics of TEs within eukaryotic genomes across evolutionary timescales and was developed to incorporate as many biologically relevant parameters into the model as possible. It uses a modular approach, beginning with a synthetic genome (Synthetic Genome Mode) or a user-provided (Custom Genome Mode) that represents a present-day or theoretical ancestral genome. Users may generate these inputs entirely from scratch with PrinTE’s built-in parameterization tools, import prior PrinTE runs for continued simulation, or use TEgenomeSimulator ^33^ outputs, which are supported by PrinTE. PrinTE implements two methods for simulation, which are the variable rate method and the fixed rate method. The variable-rate method aims to incorporate consequences from some of the biological and selective forces that guide TE evolution. We established several rules in this method: 1) The host genome survival and TE proliferation are conflicting forces pitted against each other; 2) TE superfamilies compete with each other since they occupy the same limited genomic space; 3) The rate of TE insertion is positively correlated with the number of intact TEs in the previous iteration; and 4) TE deletions are guided by the number of TEs that are deleterious to the host’s survival. For this purpose, the simulated genome consists of three components, all of which are receptive to new TE insertions, namely existing TEs, genic space, and intergenic space. If a TE inserts into another TE or intergenic space, there is no cost to the host and no impact on the TE deletion rate. In contrast, TE insertions into genic space are costly to the host, resulting in elevated TE deletion rates in the next generation. Thus, the outcome of the variable-rate method is less predictable, which may lead to generations of genome size increase followed by subsequent generations of decrease. It is useful for those who wish to explore the regulation of TE activities through time and to understand the forces that affect TEs’ bursts and purges.

The fixed-rate method is designed to generate more controlled and predictable outputs. In this method, per-generation TE insertion and deletion rates are predefined by the user, allowing the modeling of known evolutionary trajectories and the generation of results that more closely match expectations (**Fig. S1**). In practice, forward simulation should begin from a simulated genome that mirrors a present-day version and forecast a few million generations of TE evolution into the future. Alternatively, it can start from a genome as we expect it may have looked a few million generations ago and simulate it to the present day. This approach allows users to test hypotheses relating to TE bloat and purge and to potentially superimpose TE evolutionary narratives onto the species of interest.

PrinTE can simulate as briefly as a single generation, which can generate subtle changes to the simulated genome. To allow for simulating millions of generations, PrinTE employs a stepwise approach in which it can simulate TE evolution of a number of generations in steps until it reaches the total number of generations specified by the user. To approximate TE mutation in multiple generations, PrinTE uses a normalized exponential decay distribution to guide DNA substitutions, mimicking the natural turnover of TE sequences that would occur during iterative steps.

PrinTE represents a state-of-the-art simulation of TE dynamics in eukaryotic genomes, enabling the modeling of TE contributions to genome evolution and the estimation of past and future genome evolution trajectories.

#### Performance and benchmarking

The 5’ and 3’ LTR regions of an intact LTR-RT are identical upon transposition. Under the molecular clock hypothesis, mutations to the LTRs reflect the amount of time since transposition ^30,31^. Accurately estimating the sequence divergence of LTR-RTs is critical for the inference of TE activity over evolutionary epochs. We developed a k-mer-based LTR-RT dating approach, called Kmer2LTR (https://github.com/cwb14/Kmer2LTR.git), to evaluate PrinTE’s accuracy in LTR-RT modeling and its ability to forward-evolve a genome. PrinTE allows users to customize the transition/transversion ratio (Ti/Tv), which enables evaluating various nucleotide substitution models that were incorporated into PrinTE. For comparisons, we used SLiM to simulate genome divergence based on a simple neutral model. Single-nucleotide polymorphisms (SNPs) were projected onto the SLiM-simulated genealogies to obtain whole-genome sequences that were comparable to PrinTE simulations. Divergence of SLiM-simulated genomes was estimated based on mutations detected using Minimap2 whole-genome alignments.

In all comparisons, the Kimura two-parameter (K2P) model outperformed p-distance estimates, regardless of the Ti/Tv ratio (**Fig. 2A**). The one-parameter Jukes-Cantor 69 (JC69) model performed equally as well as K2P when Ti/Tv=1, but underperformed when Ti/Tv=10 (**Fig. 2A**). PrinTE and SLiM/Minimap2 each show high accuracy in their ability to detect mutations up to 25% sequence divergence, but the latter quickly dropped out at divergences exceeding 25%. PrinTE’s Kmer2LTR approach benefits from pre-determined LTR boundaries, which allow it to force a global sequence alignment onto the otherwise highly divergent LTRs and accurately detect mutation percentages up to 30%. Divergences at >30% are detectable by PrinTE’s LTR-RT approach with a slight reduction in accuracy (**Fig. 2A**). Simulations at higher generation counts lead to a wider range of sequence divergence in LTR-RTs, but the overall divergence estimate remains centered on the expected values even at the relatively high divergence range (**Fig. 2B**). These results demonstrate a robust LTR-RT divergence estimation by PrinTE’s Kmer2LTR method, especially when using the K2P model.

**Fig. 2.**
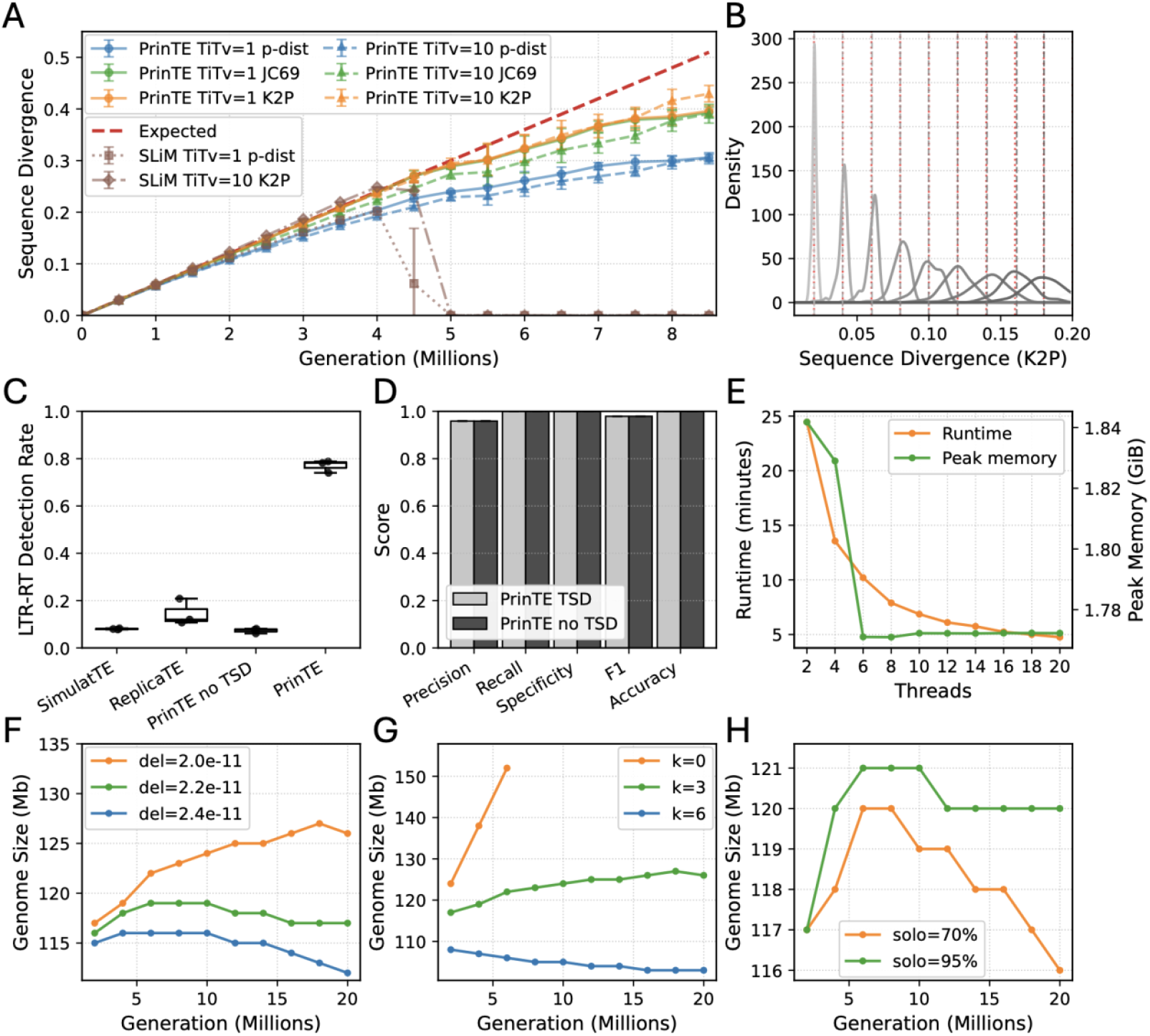
Benchmarking PrinTE. (**A**) Estimation of LTR retrotransposon divergence simulated by PrinTE with p-distance (p-dist), Jukes-Cantor 69 (JC69), and Kimura two-parameter (K2P) DNA substitution models and transition/transversion ratios (TiTv) of 1 and 10. Single-nucleotide polymorphisms were projected onto SLiM-simulated genealogies and detected using Minimap2 whole-genome alignment. (**B**) Density distribution of LTR sequence divergence simulated under various divergence scenarios and estimated by Kmer2LTR. Vertical grey dashed lines show the cumulative sequence divergence of all LTR-RTs in the simulated genome. Vertical red dashed lines show the expected sequence divergence at each time point. (**C**) LTR-RT recall rates by LTR_retriever using three TE simulators. TSD, target site duplication. (**D**) Performance metrics of LTR-RT identification by RepeatMasker using the PrinTE library and the simulated genome as input. Error bars show 95% confidence intervals (n = 1,000). (**E**) PrinTE runtime and memory utilization using different numbers of CPU threads. Genome size variation with deletion rates (**F**), deletion length preference parameter k (**G**), and solo-LTR frequencies (**H**).

Next, we assessed the simulation fidelity in PrinTE and existing TE simulators, SimulaTE and ReplicaTE. The success of structure-based TE annotation depends on the presence of known sequence structures and motifs. We reasoned that intact TEs in simulated genomes should be detectable using a structure-based annotation approach. However, existing TE simulation tools often fail to generate target site duplications (TSD), a key signature of LTR-RT integration ^30^. Our results show that LTR-RTs simulated by PrinTE have a significantly higher detection rate than those that were simulated using ReplicaTE and SimulaTE (77% vs 15% and 8%, respectively), suggesting that PrinTE was better at reflecting the biological features that compose a TE (**Fig. 2C**). We used RepeatMasker with PrinTE-derived TE library to evaluate detection accuracy of PrinTE simulations and observed very strong quality metrics (F1 = 0.978), suggesting that the output TE library and corresponding genome were accurately simulated (**Fig. 2D**).

To evaluate the computational performance of PrinTE, we simulated one million generations of TE evolution into a 100Mb genome using a step size of 100,000 generations. The initial genome was seeded with 6,000 TEs, and each subsequent iterative step resulted in insertion and deletion of ∼500 TEs. PrinTE demonstrated efficient memory usage with peak memory less than 2GB. PrinTE runtimes scale well at low thread counts and begin to saturate with increased thread allocation (**Fig. 2E**). These performance metrics demonstrate that PrinTE is highly efficient, optimized for large-scale analysis, and capable of running on personal computers or compute servers.

### Genome Size Evolution Through TE Expansion and Contraction

LTR-RTs are the primary contributing TEs to genome size in many plant, fungi, and animal genomes ^32^. To isolate the effect of TEs on genome size across temporal gradients, we focused on modeling LTR-RT evolution using PrinTE’s fixed-rate method to simulate predictable patterns of TE expansion and contraction. Here, we simulated two dynamic scenarios subsequently on the same genome. The first scenario used a TE deletion rate fourfold higher than that of insertions. We observed a 45% decrease in genome size after five million generations of forward simulation (**Fig. 3A**). The decrease was initially driven by a swift reduction in the number of intact TEs (**Fig. S2A**) followed by an increase in fragmented TEs and solo LTRs (**Fig. 3A, Fig. S2B**).

**Fig. 3.**
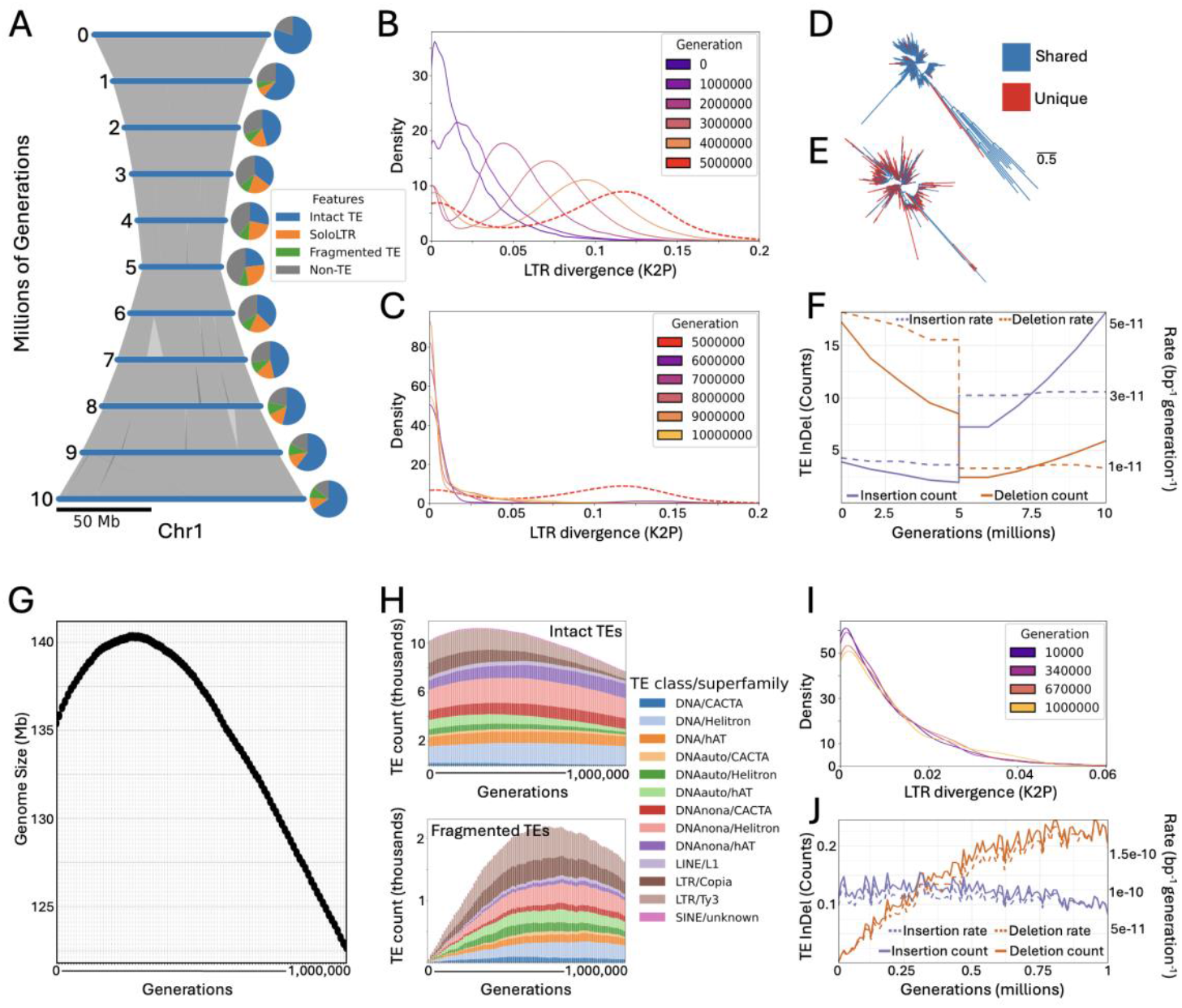
Modeling TEs’ contribution to genome size evolution using fixed (A-F) and variable (G-J) rate methods. (**A-F**) Ten million generations of LTR-RT evolution explores their long-term impact on genome size using the fixed rate method. For demonstration, we set a rate of TE deletion that exceeded insertion by fourfold in the first five million generations, then reversed the rates by setting the insertion rate exceeding deletion by fourfold over the next five million generations. (**A**) A syntenic ribbon plot illustrates the change in chromosome size. Pie charts corresponding to each time point show the relative contribution of each feature to genome size. (**B**,**C**) Density distributions of LTR-RT divergence estimated using the K2P model during the first five million generations (**B;** genome size decrease) and the second five million generations (**C;** genome size increase). Five million generations is the transition between genome size decrease and increase, which is shown in both plots as dashed red lines. Phylogenetic trees show the evolutionary relationship of LTR-RTs from the *Tekay* family after four million generations of genome size decrease (**D**) or four million generations of genome size increase (**E**). (**F**) LTR-RT insertion and deletion counts (solid lines) and rates (dashed lines) across 10 million generations. (**G**,**H**) A simulated genome replica of *Arabidopsis thaliana* is forward-evolved one million generations using PrinTE’s variable rates method. (**H**) Intact (top) and fragmented (bottom) TE counts per superfamily across one million generations. (**I**) The distribution of LTR-RT divergence estimated using the K2P model. (**J**) TE insertion and deletion counts (solid lines) and rates (dashed lines) illustrate the self-adjusting functionality of PrinTE’s variable rate method.

Following the genome size decrease, we continued the simulation but reversed the TE insertion rate to fourfold higher than that of deletions. The elevated insertion rate caused rapid genome bloating (**Fig. 3A**) powered by an increase in the numbers of both intact and fragmented TEs (**Fig. S2C**,**D**). The sequence lengths of solo, fragmented, and intact LTR-RTs shifted in response to purging pressures from the host (**Fig. S3**). Interestingly, because k>0 leads to a higher probability for deleting longer LTR-RTs, the length of intact LTR-RTs was overall reduced in the genome size decrease phase and slowly increased in the genome size increase phase when purging pressure was abated (**Fig. S3**).

During simulations of genome size decrease, when the rate of TE deletions exceeded the rate of insertion, we observed unbiased removal of LTR families through generations. Accumulation of DNA substitutions and reduced LTR-RT insertions led to elevated sequence divergence of the remaining pool of intact LTR-RTs and a shift in the age distribution across time (**Fig. 3B**). There were two distinct density peaks as simulation proceeds, even though the rate of TE insertions and deletions remains constant (**Fig. 3B**). The smaller peak represents younger LTR-RTs that were inserted recently and have had less time to be purged, while the larger peak that shifts to the right and decreases in density through time was retained from the standing body of LTR-RTs in the ancestral genome. In cases where the rate of TE insertions exceeds deletions, young LTR-RTs quickly proliferate in high abundance and dominate the density distribution of LTR-RTs in the simulated genome, masking the signal of remaining older LTR-RTs (**Fig. 3C**). We conclude that the divergence distribution of intact LTR-RTs reflects their insertion and deletion dynamics through time.

When species become reproductively isolated from each other, their TEs are also isolated and evolve independently. Genomes share TEs that derive from their common ancestor and also contain novel insertions that are unique to the other. With PrinTE, we simulated a speciation event by evolving a genome for a period of time, then diverted the genome into two isolated evolutionary paths with shifted TE dynamics to create divergent trajectories of genome expansion or contraction (**Fig. 3D,E**). In the contracting genome, there are relatively few unique TEs created after speciation (**Fig. 3D**; **Fig. S4**). In contrast, the expanding genome had a higher abundance of young and unique TEs (**Fig. 3E**; **Fig. S4**). In the fixed-rate method, the rate of TE insertion and deletion remains relatively stable, only deviating by the stochastic variability introduced by sampling a Poisson distribution, while the number of insertions and deletions introduced scales with the size of the genome (**Fig. 3F**).

Further, we profiled three deletion-related parameters in PrinTE to contextualize their impact to genome size, namely deletion rate, TE length bias, and solo LTR rate (**Fig. 2F-H**). Deletion rate controls the number of TEs that are processed by PrinTE’s deletion engine. Expectedly, we observe decreasing genome size with increasing deletion rates (**Fig. 2F**). Deletion of TEs is mainly achieved by unequal or illegitimate recombination, which both operate through sequence homology between TE sequences. In PrinTE, we allow users to set TE deletion bias exponentially to the TE length using the `-k` parameter, which can approximate the disproportionate impact of illegitimate recombination on purging longer TEs. Larger `k` has a stronger bias in deleting longer TEs and thus has more constraints on genome size (**Fig. 2G, Fig. S5**). When `k` is set to 0, TE deletion has no bias to TE length, allowing genome size growth through the accumulation of long TEs (**Fig. 2G, Fig. S5**).

Illegitimate recombination between LTR regions of individual LTR-RTs will partially remove the element and create solo LTRs as remnants. Given a fixed deletion rate, PrinTE can model the proportion of LTR deletions that go through this process by implementing the “`--solo-rate`” parameter. Increased solo rate can increase the number of solo LTRs in the genome and slow down the purging of LTR-RTs, resulting in a larger genome size (**Fig. 2H, Fig. S6, Fig. S7**). By adjusting parameters of deletion length bias and LTR solo rate that interplay with TE insertion and deletion rates, we can create complex scenarios for modeling genome size evolution in different systems.

Recognizing that the rates of TE insertion and deletion can be dynamic through time and may be modified by varying biological processes, we implemented the variable-rate method in PrinTE, which can be used to explore specific molecular features that constrain TE evolution and its impact on genome size. Here, rather than setting fixed TE insertion and deletion rates, we start with an initial rate and allow the tool to dynamically determine subsequent insertion and deletion rates based on the genetic makeup of the most recently simulated genome (**Fig. 3J**). We simulated the genome size and TE evolution of the present-day *Arabidopsis thaliana* to forecast one million generations of TE evolution using the variable-rate method in PrinTE. With the tested conditions, there is an initial period of genome bloating followed by a longer period of TE purging and genome size contraction (**Fig. 3G**). Expectedly, genome size increased through the accumulation of both intact and fragmented TEs and decreased due to the reduction of intact TEs (**Fig. 3H**). The divergence of intact LTR-RTs reflects the genome size expansion and contraction (**Fig. 3I**). These features allow us to simulate TE dynamics close to what is observed in eukaryotic genomes.

### Genome Size Evolution of Rust Fungi

We further studied the impact of TE dynamics on genome size evolution using rust fungal genomes due to their wide range of genome sizes. The *Pucciniomycotina* is a subdivision of fungi that contains 10 classes and over 8400 species ^35^. We downloaded 58 *Pucciniomycotina* fungal genomes from NCBI and JGI MycoCosm ^34^ to investigate genome expansion and contraction (**Table S1**). We identified and removed alternative haplotypes from these genomes using Redundans ^36^. The resulting haploid genome size varies from 13.6 Mb (*Mixia osmundae*) to 1018.1 Mb (*Austropuccinia psidii*), representing over 70-fold differences (**Table S2**). We performed Benchmarking Universal Single-Copy Orthologs (BUSCOs) evaluations using compleasm ^37^ and the odb12 *Basidiomycota* database. The complete BUSCOs (single and duplicated) averaged 90.7% (SD = 5.4) and contig N50 averaged 846 kbp (ranging from 111 kbp to 3.8 Mb), suggesting high assembly quality of haploid genomes (**Table S2**).

We identified the shared single-copy BUSCOs and used IQ-TREE ^38^ to construct the phylogeny of *Pucciniomycotina*. The resulting phylogeny was over-represented by two classes, *Microbotryomycetes* and *Pucciniomycetes* (**Fig. 4A**). *Microbotryomycetes* genomes are small, ranging from 17.5 Mb to 50.3 Mb. Correspondingly, on average, only 0.4% of the genome was represented by LTR sequences with a range of 0% to 3.0% (**Table S2**). This observation suggests that the best strategy for controlling genome size is to avoid the invasion of LTR-RTs. *Pucciniomycetes* have widely distributed genome sizes, ranging from 26.8 Mb to 1018.1 Mb, with an LTR sequence content corresponding to 0.4% to 67.4% (**Table S2**).

**Fig. 4.**
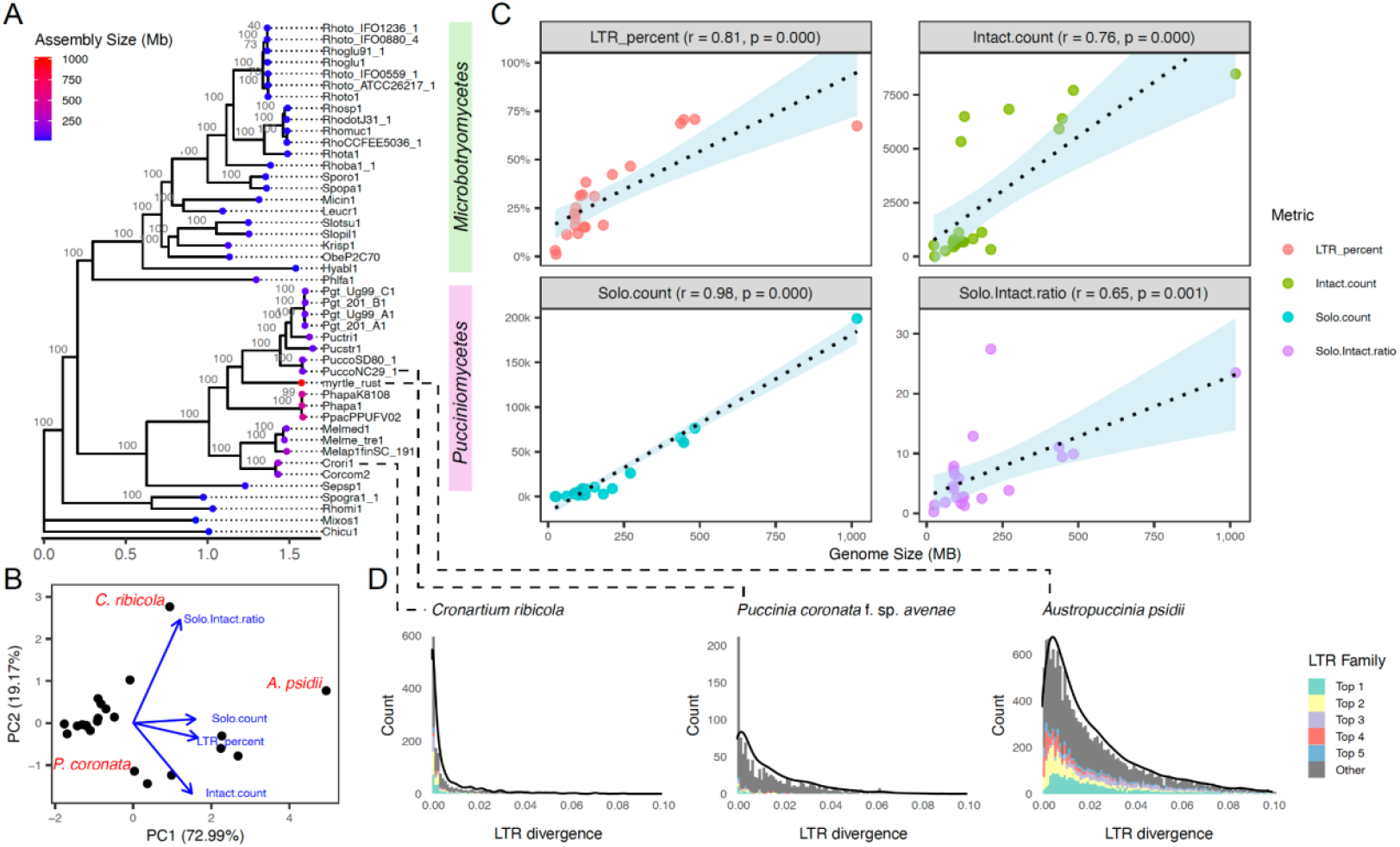
(**A**) The phylogeny of *Pucciniomycotina* fungal genomes used in this study. Protein sequences of shared single-copy BUSCO were used for tree inference with 1000 bootstrap replications. Bootstrap support percentage is shown on each branch. The two classes, *Pucciniomycotina* and *Microbotryomycetes*, are labeled beside the phylogeny. The genome size of each species is indicated by color dots. (**B**) Principal Component Analysis (PCA) on LTR metrics in fungal genomes. Each dot represents one genome. PCA loadings of each metric are shown. Genomes selected for further studies are labeled. (**C**) Pearson’s correlation (*r*) between genome size and LTR sequence content (LTR_percent), intact LTR-RT count (Intact.count), solo LTR count (Solo.count), and solo:intact ratio. Each dot represents one genome. The correlation and p-value are shown on the panel title. (**D**) Divergence of intact LTR-RTs in *Cronartium ribicola* (white pine blister rust), *Puccinia coronata* f. sp. *avenae* (oat crown rust), and *Austropuccinia psidii* (myrtle rust) genomes. The top five most abundant LTR families in each genome are colored, and they are not shared between genomes. A density line is shown as the contour in each plot. The locations of these species on the phylogeny are indicated by dotted lines.

To further study the dynamics of LTR-RTs in these fungal genomes, we *de novo* identified intact LTR elements and solo LTR sequences, and calculated the whole-genome solo:intact ratio using LTR_retriever ^30^. Using genomes with >1% LTR sequence content (n = 22), we performed Principal Component Analysis (PCA) on LTR metrics and genome size. The analysis revealed clustering among smaller genomes, whereas larger genomes appeared as outliers, including *Cronartium ribicola* (white pine blister rust), *Puccinia coronata* f. sp. *avenae* (oat crown rust), and *Austropuccinia psidii* (myrtle rust; **Fig. 4B**). PCA loadings suggested that solo LTR number, intact LTR number, and their ratio were all drivers to the outliers of large genomes (**Fig. 4B**). Including total LTR content, these LTR metrics showed strong positive correlations with genome size (Pearson’s *r* ≥ 0.65, p ≤ 0.001; **Fig. 4C**). This observation supports LTR-RTs’ major role in impacting genome size upon successful invasion.

Not surprisingly, intact LTR elements, as the product of LTR-RT proliferation, were strongly correlated with genome size (*r* = 0.76; **Fig. 4C**). However, solo LTR sequences as the product of LTR-RT removal ^30^ showed an even stronger correlation, almost linearly correlated with genome size (*r* = 0.98; **Fig. 4C**). Another counter-intuitive observation is the positive correlation between genome size and solo:intact LTR ratio, the latter is a metric used to measure the intensity of LTR-RT purging ^31^. This observation contradicts the conventional understanding that the higher the solo:intact LTR ratio, the more intense purging of LTR sequences, and thus the smaller the genome ^39^. The positive correlation between solo:intact ratio and genome size suggests that illegitimate recombination alone may not be an effective way of escaping genome obesity.

We selected *C. ribicola, P. coronata* f. sp. *avenae*, and *A. psidii* as representatives of three distinct roadmaps to genome size evolution. *C. ribicola* has a genome size of 211.6 Mb with the LTR divergence distribution showing an extremely quick turnaround time for LTR element insertions (**Fig. 4D**; **Fig. S8**). The solo:intact ratio of *C. ribicola* is 27.4 (**Table S2**), the highest among all genomes analyzed in this study, suggesting a quick insertion–quick removal mode of action, and a relatively stable or slowly increasing genome size. The *P. coronata* f. sp. *avenae* has a genome size of 113.2 Mb and a solo:intact ratio of 1.7 (**Table S2**). The LTR divergence distribution showed a much slower turnover time for LTR-RT insertions (**Fig. 4D**; **Fig. S8**), suggesting a slow insertion–slow removal mode of action, and a relatively stable genome size. *A. psidii* has an extremely large genome size of 1018.1 Mb and a solo:intact ratio of 23.5, the second highest ratio among our data (**Table S2**). However, the LTR divergence of *A. psidii* shows the peak of LTR-RT insertion passed the current time (**Fig. 4D**; **Fig. S8**), suggesting a slowdown of insertion activities ^8^ and a decreasing genome size.

### Simulating Digital Replicas of Rust Fungal Genomes

TE composition in real genomes varies substantially across TE families and species because of their complex and diverse evolutionary histories. This variability makes it challenging to simulate genome-wide TE landscapes that can be used for modeling and hypothesis testing. To address this, we used TEgenomeSimulator ^33^ with its TE Composition Approximation Mode to model the TE profiles of *C. ribicola, P. coronata* f. sp. *avenae*, and *A. psidii* genomes within the *Pucciniomycotina* subdivision (**Fig. 5, Fig. S9, Fig. S10**). The sizes of these simulated genomes closely matched their actual genome sizes. Similarly, the TE proportion of the simulated genomes closely approximated the observed values in terms of TE loci and TE bases per superfamily (**Fig. 5AB, Fig. S9, Fig. S10**). TEgenomeSimulator estimated the sequence divergence and integrity distributions of each TE family in the real genomes for the simulation of digital replicas. The simulator uses normal distributions to smooth out small-scale fluctuations present in the real genome, resulting in a simulated divergence profile that broadly resembled the real data (**Fig. 5C, Fig. S9, Fig. S10**).

**Fig. 5.**
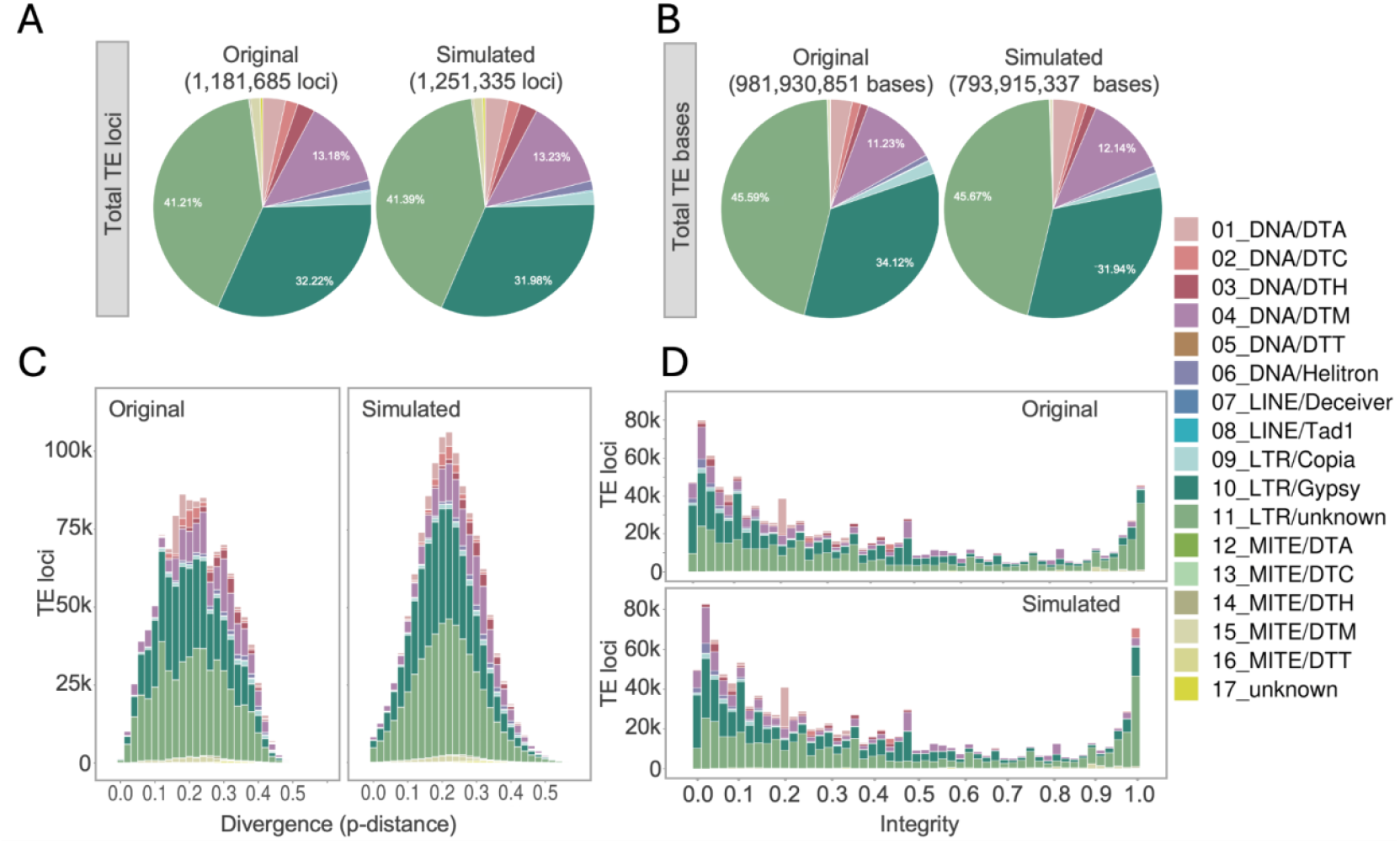
TE Composition Approximation for the *Austropuccinia psidii* genome. (**A**) Comparison of total TE loci between the original and simulated genomes. (**B**) Comparison of total TE bases between original and simulated genomes. (**C**) TE sequence diversity distribution colored by TE superfamily. (**D**) The distribution of TE sequence integrity, which is the length ratio relative to the consensus sequence, is colored by TE superfamily as in (**C**).

TEs can break down over time due to nested insertions, recombination, small deletions, and other processes. The whole-genome distribution of TE integrity clearly reflects the turnover rate of TEs in the three selected fungal genomes (**Fig. 5D, Fig. S9D, Fig. S10D**). The *C. ribicola* genome contains mostly high-integrity TEs, suggesting a high rate of TE turnover and active TE insertions (**Fig. S9D**). In contrast, the *P. coronata* f. sp. *avenae* genome harbors highly fragmented TEs and very few intact TEs, suggesting a low activity profile (**Fig. S10D**). The myrtle rust genome, *A. psidii*, presents a bimodal distribution in TE integrity, suggesting both heavy bloating and purging may have been prominent drivers in the evolutionary history of the genome (**Fig. 5**). The digital replicas of these genomes have distributions of TE integrity that are almost identical to the real genomes (**Fig. 5D, Fig. S9, Fig. S10**). These simulations yield generalized distribution patterns that capture the major trends in TE sequence evolution of each genome, providing a robust foundation for PrinTE’s simulations of TE bloating or purging episodes, effectively serving as burn-in phases for modelling genome evolution.

### Accessing Genome Evolution with Forward Simulations of LTR Activities

Our simulations using PrinTE present an effective capturing of LTR-RT dynamics using the divergence distribution (**Fig. 2A-C**), which, in reverse, can be used to empirically estimate parameters of LTR-RT dynamics. Here, we demonstrate the utility of PrinTE in modeling genome size evolution under the impact of TE bloat and purge. Despite their shared ancestry (**Fig. 4A**), the three selected fungal species (*C. ribicola, P. coronata* f. sp. *avenae*, and *A. psidii*) show immense variation in genome size and LTR-RT evolution. To test our hypothesis that *C. ribicola* has a stable genome size that it achieved by rapid but balanced LTR-RT insertion and deletion, we forward-evolved the *C. ribicola* genome digital replica using PrinTE with a variety of insertion and deletion rates in search of an LTR-RT divergence distribution that closely matches the observed distribution. At each trial, we compared the LTR-RT divergence distribution with that of the real genome (**Fig. S11)**. We found that an LTR-RT insertion rate of 2×10^-9^ per bp per generation and a deletion rate of 2.4×10^-9^ per bp per generation provided a simulated genome that closely matched the *C. ribicola* genome, with the genome size increased moderately through time (**Fig. 6A,J**). Forward-projecting 200,000 generations of TE evolution indicated that the density distribution of LTR-RT ages in the simulation closely matched that of the real genome (**Fig. 6A, Fig. S9**). The solo:intact ratio of the simulated genome was 13 (**Fig. 6C**) in comparison to 27.4 in the real genome (**Table S2**). Although the forward-simulated genome does not share an identical LTR-RT footprint to that of *C. ribicola*, its proximity to the real data suggests that PrinTE can effectively infer TE dynamics through forward simulations. Simulation of the *C. ribicola* genome suggests that the species might have inserted and deleted (either intact or fragmented) one effective LTR-RT every two generations during the last 200,000 generations.

**Fig. 6.**
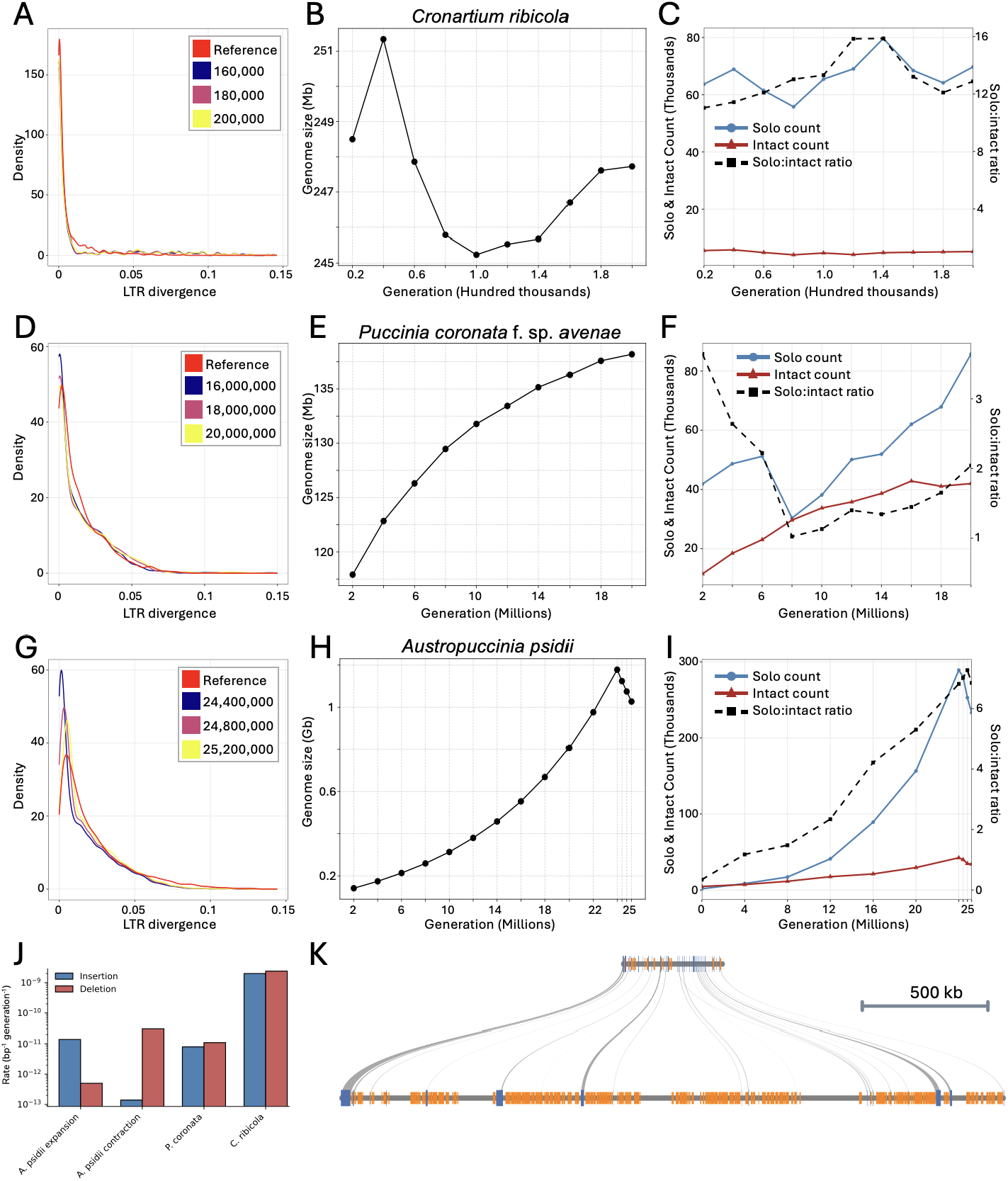
Simulating genome size evolution in three *Pucciniomycetes* genomes. (**A-C**) *Cronartium ribicola* (white pine blister rust), (**D-F**) *Puccinia coronata* f. sp. *avenae* (oat crown rust), and (**G-I**) *Austropuccinia psidii* (myrtle rust). (**A, D, G**) Sequence divergence compares the empirical divergence distribution of LTR-RTs (Reference) to simulated genomes. LTR divergence was estimated using LTR_retriever with the K2P model. Three timepoints were plotted for *C. ribicola* (**A**) and *P. coronata* f. sp. *avenae* (**D**) to showcase the relatively stable TE dynamic. The three timepoints of *A. psidii* (**G**) highlight the rapid purging of LTR-RTs during the period of genome size contraction. (**B, E, H**) Genome size changes across generations. (**C, F, I**) The empirical solo:intact ratio estimated using LTR_retriever. (**J**) The estimated LTR-RT insertion and deletion rates of fungal genomes. The expansion and contraction phases of the *A. psidii* simulation were plotted separately. (**K**) Microsynteny between the simulated *P. coronata* f. sp. *avenae* before (top) and after (bottom) the TE evolution that led to genome size increase in the simulated *A. psidii* genome. Collinear genes are blue, and LTR-RTs are orange.

We repeated the same process using the genomic digital replica of *P. coronata* f. sp. *avenae*a to test our hypothesis that the species has slow TE insertion and slow removal, leading to a relatively stable genome size. An LTR-RT insertion rate of 8×10^-12^ per bp per generation and a deletion rate of 1.10×10^-11^ per bp per generation were found to be optimally adapted for maintaining the LTR-RT characteristics of the real genome for over 20 million generations (**Fig. 6B,J**; **Fig. S12**). Dating LTR-RTs in the forward-simulated and the real genome showed that the distributions were nearly identical (**Fig. 6D**). The solo:intact ratio in the forward-simulated genome mimics that of the real genome, at 2.0 and 1.7, respectively (**Fig. 6F, Table S2**). With these rates, we see a gradual increase in average genome size, adding approximately 1.25 Mb per million generations (**Fig. 6E**). These results suggest that, in contrast to *C. ribicola, P. coronata* f. sp. *avenae*a appears to have an almost inactive LTR-RT footprint with an average of 1090 generations between each effective LTR-RT insertion and an effective deletion every 790 generations.

For our final simulation, we used PrinTE to create an initial burn-in genome that mimicked the gene and TE landscape of a hypothetical ancestral genome of *A. psidii*, highlighting PrinTE’s utility as a stand-alone TE simulation and evolution tool. We began with a relatively small genome of 110Mb that is typical in the *Pucciniomycetes*, and simulated the TE evolution that led to the enormous genome bloating of *A. psidii*. Based on the genome size and LTR-RT divergence distribution (**Fig. 4A,D**), we expected that TE insertions exceeded deletions for a long period of time, leading to genome size increase, followed by a shorter period of rapid LTR-RT deletions. Through evaluation of various dynamic parameters, we found that an insertion rate of 1.4×10^-11^ per bp per generation and a deletion rate of 5×10^-13^ per bp per generation for 24 million generations was optimally tuned to mimic the observed LTR-RT age distribution of the *A. psidii* genome during the long period of genome size increase (**Fig. 6G,J**; **Fig. S13A**,**B**). During this period, we aimed to achieve a genome size that was marginally larger than the real genome so that we could tune up the rate of deletion to reflect the more recent decline in young LTR-RTs observed in *A. psidii* (**Fig. 4D**). At peak LTR-RT invasion, the simulated genome was 1,178 Mb in size (**Fig. 6H**).

The drop of recent LTR-RT insertions in *A. psidii* suggests a reduced insertion rate and an increased deletion rate. Using a substitution rate of 2.0×10^-9^ per bp per generation ^40^, we estimated that the insertion reduction phase of *A. psidii* had lasted for 1.2 million generations. We then forward-simulated an additional 1.2 million generations using elevated deletion rates on top of the increasing phase of simulation. Systematic evaluation of TE dynamic parameters revealed that an insertion rate of 1.4×10^-13^ per bp per generation and a deletion rate of 3×10^-11^ per bp per generation can achieve a genome size that is near *A. psidii*, at 1,027 Mb (**Fig. 6H,J**; **Fig. S13C**), and the LTR-RT divergence distribution mimics the real genome (**Fig. 6G**; **Fig. S13D**). The solo:intact ratio of the simulated *A. psidii* genome terminated at 6.8 (**Fig. 6I**), which is lower than the observed value at 23.5 (**Table S2**), suggesting other evolutionary processes also play roles in shaping the modern-day *A. psidii* genome.

Simulation of *A. psidii* suggests two major phases that contributed to the present-day genome size. The first was a long period of genome size increase in which LTR-RT insertions exceeded deletions by 28-fold, and the second was a short period where deletions exceeded insertions by 214-fold. Our results support a recent contraction model for genome size evolution in *A. psidii* (**Fig. 6J**). It is important to note that the estimated parameters are effective insertion and deletion rates averaged from a long period of evolutionary time, which may not capture short periods of TE bloat and purge.

Because these three species were selected to represent the extremes of LTR-RT dynamics in the sampled *Pucciniomycetes* genomes, we suggest that the amplification and degradation of LTR-RT can vary widely within the fungal kingdom. In future studies, it will be interesting to see if such a wide variety of LTR-RT dynamics exists at narrower phylogenetic levels.

## Discussion

The TE composition profile of a genome serves as an evolutionary archive, preserving signatures of TE dynamics and host genome responses that collectively shape genome evolution. Deciphering these patterns from empirical data is challenging. Whether to test evolutionary hypotheses or to predict future trajectories, simulation is often necessary. To address this, we built PrinTE, a flexible, forward simulation framework for transposon evolution. PrinTE significantly expanded the capacity for simulating complex TE dynamics and testing evolutionary hypotheses.

PrinTE allows forward simulation on TE dynamics

Transposable elements are a major genomic component and subject to all evolutionary forces, including purifying selection, drift, and mutation. The ability to trace TEs’ responses to these events will greatly facilitate the studies of TE dynamics. Forward simulation is a key approach that allows the artificial evolution of TEs and other genomic features, which provides opportunities to sample the intermediate state of the evolutionary process.

SLiM is a powerful forward simulation program that can simulate the majority of evolutionary scenarios ^24^. However, its operation on single-nucleotide variants limits its application on TEs with varying lengths and abundance of nestings. SLiM provides a workaround by first compressing the whole TE locus into a presence and absence unit and performing simulations on these units. Sequence content is then filled back to replace the units to generate genomic sequences. This approach makes it difficult to model the variation accumulated through generations on the simulated TE sequences, including the deletion and formation of solo LTRs, which are critical for genome evolution studies. On the contrary, PrinTE operates on whole-genome sequences, including invariant sites that allow intuitive modeling of complex scenarios and interpretation of results. In particular, PrinTE models the insertion and deletion rates of TEs and generates solo LTRs, which empowers the study of genome size evolution driven by TE bloat and purge.

The input files of PrinTE are highly flexible (**Fig. 1**). It has a built-in engine to simulate highly customizable starting genomes, in which users can specify genome size, TE content, TE superfamily abundance, and TE divergence in their simulations. PrinTE integrates CDS sequences in the simulation, allowing for modeling purifying selections, tracing large structural variants, and studying the evolution of gene space. PrinTE also takes simulated genomes from TEgenomeSimulator, allowing forward simulation from different diverse backgrounds of genomes. Most importantly, PrinTE can take its own output as input (e.g., **Fig. 3A-F**), allowing for the simulation of varying TE dynamics and studying the host-TE arms race. This feature also enables the simulation of genomic divergence and speciation driven by differences in TE activities.

### TE contributes to genome size expansion

The contribution of TE contents to genome size expansion has been confirmed across many taxa and in our data (**Fig. 4C**). Over time, genomic data have come faster and of a higher quality, and yet, as noted by Osmanski *et al*. (2023) ^41^, the role of TEs in genome architecture and evolution is often an afterthought. More in-depth analyses are restricted to domain specialists for a meaningful evolutionary perspective.

In order to put forward mechanistic hypotheses, authors have developed specialized computational methods to detect signals of TE radiation and impacts on speciation and genome size. For example, Ricci *et al*. (2018) ^13^ developed a measure of TE Density of Insertion and Relative Rate of Speciation to model TE radiation bursts and silencing. Kapusta *et al*. (2017) ^12^ estimated TE-driven DNA gains and losses and proposed the “accordion” model of TE radiation followed by DNA loss for tight genome size maintenance in mammals and birds. Osmanski *et al*. (2023) ^41^ applied a Redundancy Analysis to study TE composition across mammalian taxa generated through the Zoonomia genomes ^42^, and found evidence of multiple recent TE expansions driving larger genome sizes and wide-spreading TE-silencing events across the mammalian tree. These authors hypothesize that variations in TE defense mechanisms may have played a role in whether specific lineages were overcome by a TE burst at a given evolutionary time.

However, we have limited knowledge about the maintenance of compact genomes despite evidence of TE radiation bursts and the maintenance and regulation of extremely large genomes. The Chinese pine attributes 69.4% of its 25.4 Gb genome to TEs, 86.5% of which are made up of LTR-RTs ^16^. The Chinese pine genome showed no sign of recent whole-genome duplication, but 91.2% of its genes were undergoing discrete duplication or family expansion, strongly associated with TEs. Our results revealed two different genome dynamics in the *Pucciniomycotina* subdivision of fungi, including the *Microbotryomycetes* class that harbours compact genome size (18-50 Mb) and the *Pucciniomycetes* class with drastic differences in genome size (27-1018 Mb) (**Fig. 4A**; **Table S2**). The *Microbotryomycetes* class is almost devoid of LTR sequences (**Table S2**), suggesting that the most effective way of maintaining compact genome sizes is to be immune to the invasion of LTR-RTs. Genomes in the *Pucciniomycetes* class have varying richness of LTR-RTs (**Table S2**), and the PCA reveals several types of LTR dynamics (**Fig. 4B**) that are likely major drivers of genome size variation.

The myrtle rust fungus expanded its genome size from an approximate ancestral size of 110 Mb to the current 1,018 Mb, experiencing a ∼10× increase. Our simulation studies revealed that the genome is experiencing a downsizing effect, with LTR-RT deletions being >200× faster than insertions (**Fig. 6**). Using PrinTE’s built-in genome simulation utility, we were able to simulate a genome that mimicked the TE landscape of the ancestral genome of myrtle rust, and further used PrinTE to simulate two phases of evolution, including a long expansion period and a short contraction period. This model fits well with the observed composition of the myrtle rust genome, suggesting a close approximation of its trajectory of genome size evolution.

In this study, we used TEgenomeSimulator and PrinTE to smooth out small-scale TE activities, then used PrinTE to simulate the real-time mobilization and deletion behavior of TEs to capture the most dominant expansion or contraction events. We recognize that TE activity is tightly linked to host genome regulation, particularly when it impacts gene function and organismal fitness. It is also important to account for more complex patterns in sequence identity distributions, which may go beyond simple normal distributions. The combined use of PrinTE and TEgenomeSimulator greatly enhanced the simulation capacity, enabling richer and more flexible hypothesis testing across both evolutionary and mechanistic contexts. Future development would benefit from incorporating features such as customizable identity distributions, more functional genomic context, epigenetic regulation mechanisms, and additional insertion site preferences.

### Conclusions

In conclusion, PrinTE is a versatile tool that allows flexible forward simulation of whole-genome sequences with fine-grained user control of genome size, TE content, superfamily abundance, TE divergence profile, TE insertion, TE deletion, TE mutation, solo LTR formation, genome divergence, and speciation. We demonstrated the use of PrinTE in resolving the genome size evolution of *Pucciniomycotina* genomes. In particular, we inferred the TE dynamic parameters underlying the genome size evolution of myrtle rust (*Austropuccinia psidii*) and other large fungal genomes. Our work transforms the study of genome evolution into a highly traceable and explorative simulation framework, and allows testing of data-driven hypotheses.

## Methods

### Implementation of PrinTE

#### Burn-in

PrinTE simulates the forward evolution of genomes. The starting point for simulation is a burn-in genome in FASTA format and a corresponding PrinTE-formatted BED file. These will serve as the starting point (0 generation) for which no forward evolution has yet taken place. The burn-in FASTA and BED can be generated *de novo* by PrinTE, or previous PrinTE outputs feeding back into PrinTE for additional forward simulations. PrinTE *de novo* burn-in allows many customizations, including genome size, chromosome number, percentage of the genome to be genic and TEs, and the ratio of each TE superfamily in the genome. Each TE insertion creates a TSD with the length determined by the TE superfamily, which was adopted from the TEgenomeSimulator codebase. In some instances, it may be beneficial to apply baseline mutation to TEs in the burn-in such that they do not exactly match elements in the TE library. This is particularly relevant for self-dating elements, like LTR-RTs, where the default TE library only contains very young LTR-RTs, with exactly identical LTRs. For baseline mutations in the burn-in genome, we apply a normalized exponential decay distribution to skew mutations heavily toward lower mutation values:

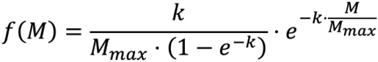

The probability density function is aimed at flexibly and accurately modeling TE age distributions as they are observed in real data, where aggressive TE purging leads to higher probability of observing young TEs (**Fig. 4D**). The parameters *M*_*max*_ (maximum mutation percentage; ‘`--TE_mut_Mmax`’) and *k* (decay constant; ‘`--TE_mut_k`’) are customizable, allowing users to adjust the shape of the distribution to reflect a wide range of evolutionary and biological scenarios and to match the observed TE half-life of any species (see **Fig. S1A**,**B**). The half-point of sequence divergence among the simulated TE is:

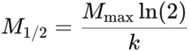

#### Two Simulation Modules

Forward simulation begins once the burn-in genome is established. Users can specify the total number of generations (‘`--generation_end`’) and a smaller number of generations for each iteration (‘`--step`’). There are three sequential modules for forward simulation: 1) genome mutation, 2) TE insertion, and 3) TE deletion. The pipeline will iterate over these processes for each step until the target number of generations is simulated. PrinTE can iterate through modules using constant and **fixed** TE insertion and deletion rates, or it can iterate using self-adjusting **variable** TE insertion and deletion rates. The fixed and variable methods begin identically, starting with the mutation of the genome. The expected number of DNA mutations per sequence (*λm*) is calculated as:

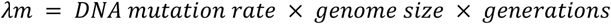

Because TE insertions can occur at any generation of a defined evolutionary window (*i*.*e*., a simulation step), they could accumulate DNA mutations less than *λm*. To capture this dynamic, we used a normalized exponential decay function similar to the decay function used for the burn-in genome to weight TE mutations toward lower values. We anchored the mutation distribution here to approximate the complicated evolutionary dynamics of TE activities. The actual number of mutations for each newly simulated TE is sampled from the probability density function, *f*(*mTE*), to help weight their observed mutations toward lower frequencies:

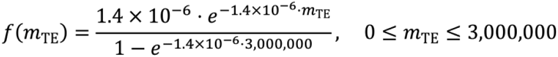

The half-life of simulated TE insertions is t_1/2_ = ln(2)/k, where k = 1.4x10^-6^, thus t_1/2_ ≈ 495,000, capturing the observation of TE half-life reported in multiple eukaryotic organisms ^30^.

#### Simulation with Fixed Rates

##### TE insertion

After DNA mutations, TEs are inserted into the genome at random locations. They may insert into intergenic regions, into other TEs, resulting in nested TE insertions, or into genes, resulting in fragmented genes. Information about nesting and fragmentation is recorded in the BED file. In the fixed rate method, the number of TEs to be inserted (*λ*_*i*−*fix*_) is calculated as:

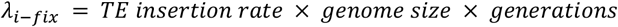

##### TE deletion and solo LTR

Donor TEs are randomly selected from the TE library based on user-defined abundance ratios per superfamily. A nested insertion will fragment the acceptor TE into two fragments. For example, if TE1 inserts into TE2, it will generate three sequences for further operations (e.g., deletion), including two resulting fragments of TE2 and the complete sequence of TE1. This differs from simple insertions into intergenic regions, where each insertion results in only one sequence for further operations. During the deletion process, full-length TEs and TE fragments are all eligible for deletion. The number of TE deletions (*λ*_*d*−*fix*_) is calculated as follows:

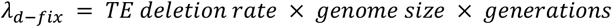

Illegitimate recombination is a prevalent process for TE deletions and is facilitated by sequence similarity between nearby homologous sequences ^31^. Longer sequences have a higher probability of causing illegitimate recombination than shorter sequences because there are more similar base pairs to initiate pairing. As a result of this phenomenon, longer TEs have a higher probability of facilitating illegitimate recombination and a higher likelihood of being deleted than shorter TEs. We used an exponential decay function to model an exponentially increasing probability of TE deletion as TE length increases:

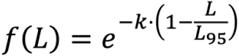

Here, *L*_*95*_ represents the TE length at the 95^th^ percentile of the length distribution in the genome, *L* represents the length of a TE (whether intact or fragmented), and *k* is a user-defined parameter (‘`--k`’) that controls the rate at which deletion probability increases with sequence length. This modification allows us to favor the deletion of longer TE features by assigning higher deletion weights to sequences that are closer in length to the longest observed TE. Except for LTR-RTs, TE deletions span the full length of the TE feature. If a nested TE is selected for deletion, the adjacent TE fragments it disrupts are reconstituted into an intact TE. LTR-RTs are an exception to full-length deletions because they can undergo recombination with themselves owing to the sequence homology between their long terminal repeats, resulting in the formation of a solo LTR. PrinTE supports deletions that lead to solo LTR formation. If an LTR-RT is selected for deletion, it will be converted to a solo LTR at a user-specified frequency, which is set by default to 95% (‘`--solo_rate`’).

##### Simulation with Variable Rates

In addition to the fixed-rate TE insertion/deletion method outlined above, in which the number of insertions and deletions per generation is constant and modeled by user-selected rates, PrinTE also offers a variable method that self-adjusts the number of insertions and deletions per generation based on genomic landscapes. This aims to replicate a subset of the biological forces that affect TE insertions and deletions. The core processes, 1) genome mutation, 2) TE insertion, and 3) TE deletion, are identical between methods. The genome mutation step is identical between methods and is described above.

##### TE insertions

In real genomes, the rate of TE transposition is influenced by the number of intact TEs, since only intact TEs are capable of initiating new insertions. Genomes with a high number of intact TEs tend to experience more transpositions per generation than those with few. Fragmented TEs and solo LTRs are no longer viable for transposition. In the variable method, the number of TE insertions (*λ*_*i*−*var*_) is modeled as follows:

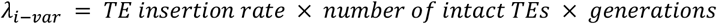

This simulates insertions via transposition, but some TEs will be acquired horizontally and drift into the genome. We set that the frequency of horizontally acquired transpositions is influenced by TE prevalence within the species. We set that species with TE-rich genomes are more likely to acquire additional TEs horizontally, while species with TE-poor genomes are less likely to do so. For this calculation, the TE composition of the genome is surveyed before simulations and used as a proxy for TE abundance in the species. horizontally acquired TEs (*λ*_*i*−*hor*_) are then simulated accordingly:

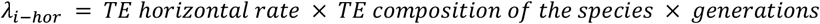

##### Insertion bias

The distribution of TEs across chromosomes is uneven and can be influenced by chromatin states, with euchromatin generally more accessible for TE insertion. In the variable method, we model this by treating genes and their surrounding regions as euchromatin, and the gene-poor genomic backbone as heterochromatin. We can modulate insertion location probability by chromatin type using a bias factor (‘`--chromatin_bias_insert`’). We assign insertion probability weights to euchromatic and heterochromatic sequences as follows:

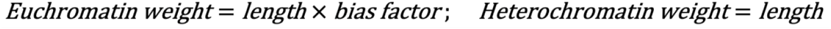

It is important to note that PrinTE does not use genes as functional units, but rather as representative segments of the genome to which cellular and selective attributes can be assigned. This enables the modeling of global genome features. Unlike the fixed-rate method, where TEs are inserted at random in proportions defined by the user, the variable-rate method inserts TEs in proportions that reflect the current genomic composition.

##### Fitness-dependent TE deletions

TEs can trigger extensive and rapid proliferation, resulting in a sudden increase in their genomic abundance. If left unchecked, this proliferation can destabilize the genome and impose a selective disadvantage on the host organism. Under the variable-rate method, the number of TE deletions is dynamically adjusted according to the extent of gene disruption caused by TE insertions. In this context, genes are modeled as sequences subject to purifying selection, whereby insertions of TEs into these regions are deleterious and thus reduce the observed number of TE insertions. The number of TE deletions (*λ*_*d*−*var*_) is then determined by the following expression:

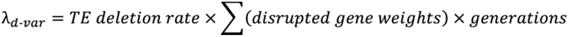

In actual genomes, not all genes that confer a selective advantage exert the same influence on fitness. While most TE insertions into genes are deleterious, some have more severe effects. To capture this variability, we model the distribution of TE selection pressures across genes using a log-normal distribution (**Fig. S1C**):

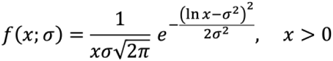

In this framework, each disrupted gene contributes a positive value to the total disrupted gene weight. The mode of the distribution is fixed at 1 to gain better control of the spread. Because of the long tail of the distribution, a small subset of genes exerts a disproportionately large influence on fitness. Consequently, TE insertions in these regions lead to a substantial increase in deletion events. The parameter σ, which is set via the command line (‘`--sigma`’), determines the spread of the distribution. Larger values of σ produce longer tails and increase the range of selective effects among genes. In reality, TE insertions in some genomic regions can be lethal. PrinTE does not simulate lethality explicitly. Instead, we assume that simulating many generations using a stepwise approach effectively filters out lethal insertions. Over time, such deleterious types are eliminated from the population and do not persist in the simulated genomic landscape.

##### TE deletion bias

Once the number of deletions is calculated, the specific TEs to delete are determined by the same exponential growth function (*f*(*L*)) used in the fixed-rate method multiplied by additional genomic attributes that might affect the probability of TE deletion. These genomic attributes include the selection coefficient parameter (‘`--sel_coeff`’) and the chromatin bias parameter (‘`--chromatin_bias_delete`’). The selection coefficient parameter is used to assign additional deletion weight toward the TEs that disrupt genes. Similarly, we can assign additional weight toward deletions in simulated euchromatin regions. Longer TEs and TEs nearer to or within impacted genes are more likely to be deleted based on user-specified thresholds.

##### LTR-RT dating

PrinTE implements a novel and accurate (**Fig. 2A,B**) approach called Kmer2LTR to estimate sequence divergence between the LTR regions of LTR-RTs. Optionally, Kmer2LTRuses known LTR region boundaries as input to initiate the LTR-RT dating process. For empirical data where the location of LTRs is unknown, we developed a k-mer approach to estimate LTR region boundaries. The k-mer approach first uses Jellyfish ^43^ to identify and count 8-12 mers. K-mers occurring exactly twice in the LTR-RT are mapped back to the LTR-RT. The resulting k-mer pairs are sorted numerically by position. Pairs that disrupted the monotonic order or that deviated by more than two standard deviations from the mean distribution of map position are removed. The map coordinates of the remaining k-mer pairs are used to establish the putative LTR region. A 65bp extension is applied to each of the putative kmer-determined LTR regions to ensure that the LTRs are fully captured. MAFFT ^44^ is used to align LTR regions within each LTR-RT, and trimAl ^44,45^ is used to prune overhanging extensions and to remove small gaps. This approach accurately identifies the boundaries of LTR regions from young to very old LTR-RTs. The gap-free and extension-cleaned LTRs are then fed to the wavefront alignment (WFA) algorithm ^44–46^ for global sequence alignment with a two-piece gap-affine scoring model to estimate sequence divergence. With these implementations, Kmer2LTR only requires a fasta of LTR-RTs and identifies the LTR regions *de novo*. This pipeline is freely available on GitHub (https://github.com/cwb14/Kmer2LTR.git).

### Searching for TE Dynamic Parameters using PrinTE

To identify optimal TE dynamics in a given genome, we employed a systematic parameter sweep approach using PrinTE. This method involved iteratively testing combinations of key parameters to approximate empirical TE behavior and genome size observed in the genome of interest. The ‘`pipeline.report`’ file contains an updated log of TE insertion and deletion counts per simulated iteration and can also be useful in monitoring PrinTE runs.

#### Parameter Space Exploration

We conducted a grid search over two critical parameters, insertion and deletion rates, governing TE insertion dynamics. Users can specify parameter ranges (upper and lower limits) and increments of parameters. Based on the hypothesis of genome size dynamics, constraints on the deletion rate can be applied to narrow down the search space. For example, if a genome evolves with a stable size, a deletion rate between 1.2 to 1.5 folds of the insertion rate can be used. We also monitor the size of simulated genomes and abort simulations when over or under the user-set size to ensure strict resource consumption. For example, if the target genome size is 150 Mb, simulation parameters that create genomes over 180 Mb in any step are deemed unfit for the hypothesis and thus aborted to reduce the computation load. PrinTE’s built-in ‘`--max_size`’ parameter can be used to set the upper limit of genome size and stop the simulation when exceeding the specified size.

Additional parameters included length bias for deletions (‘`--k`’) and solo LTR rates (‘`--solo_rate`’). In practice, we found k values of 0, 3, or 6 and solo rates from 70% to 95% (with 5% increments) could provide sufficient dynamics on top of the insertion and deletion rates. Higher k value will lead to slower genome size increase due to the preference for deleting longer TEs (**Fig. S5**). Higher solo rate will lead to faster genome size increase due to the creation of solo LTRs instead of full deletions (**Figs. S6, S7**) and increased solo:intact ratio.

#### High-Throughput Simulation Framework

PrinTE simulations were executed through an HPC cluster using array jobs (SLURM workload manager), enabling parallel processing of parameter sets. Users can set the total number of generations (‘`-ge`’) with sampling steps (‘`-st`’) for all simulations during parameter sweeps.

The species-specific substitution rate (‘`--mutation_rate`’) is also critical to the simulation outcome. In cases where an accurate substitution rate is not available for the species, users can try rates in different magnitudes (e.g., 1×10^− 8^, 1×10− ^9^, or 1×10− ^10^ per bp per generation) in the simulation, then compare the LTR divergence of simulated genomes to the real genome. Higher mutation rates will introduce more mutations to TEs in a given timeframe, thus creating gentler slopes of LTR divergence. In reverse, lower mutation rates will create steeper slopes of LTR divergence.

#### Empirical Estimation of LTR Divergence and solo:intact Ratio

Outputs were processed through a standardized pipeline to assess TE dynamics, including genome size change, LTR divergence distributions, and solo:intact ratio. Intact LTR elements were extracted from each simulation using the “`extract_intact_LTR.py`” script; their divergences were estimated with the K2P model and a mutation rate of 2.0×10^-9^ per bp per generation using the “`seq_divergence.py`” script, and divergences were plotted using the “`ltr_dens.py`” script, all available in PrinTE.

Although the BED file output of PrinTE contains the annotation of all intact and solo LTR sequences, we perform a *de novo* scan of intact LTR-RTs using LTR_retriever (v3.0.1) ^30^ on the simulated genomes, so that these results are comparable to those obtained from the real genome. The resulting LTR library was used to annotate whole-genome LTR sequences with RepeatMasker (v4.1.7-p1) ^47^. Solo LTRs were then identified by the “`solo_intact_ratio_wrap.sh`” script available in the LTR_retriever package, which also calculated the overall solo:intact ratio of each genome. For the same reason, LTR divergence distributions of simulated genomes were also plotted using intact LTR-RTs identified by LTR_retriever. The “`plot_divergence.R`” script available in PrinTE was used to plot the density of LTR divergence in simulated and real genomes. Parameter sets were ranked by their ability to reproduce three empirical features: 1) Simulated genome sizes are consistent with hypothesis predictions; 2) Genome-wide LTR divergence distributions are consistent with empirical whole-genome data; 3) Solo:intact LTR ratios are close to the empirical value. This approach identified optimal parameters where simulated TE dynamics showed convergence with multiple empirical metrics while maintaining computational feasibility.

### Benchmarking and performance testing

For benchmarking purposes, we disabled the probability density function that controls the number of LTR-RT mutations in the burnin (*f*(*mTE*)) so that LTR-RTs had mutation rates mirroring those of the genome. This allowed us to compare with other simulators that do not model such phenomena. We used PrinTE with parameters “`-P 10 -itn 150 -cn 1 -sz 100Mb -ge 7500000 -st 500000 -t 20 -k 0 -kt -m 3e-8 -sr 0 -tmx 0 -- TE_lib ./PrinTE/athrep.ltr.full.gz`”. We used extremely low rates of TE insertion and deletion (“`-F 3.0e-200,5e-200`”) to effectively disable transpositions and purging so that the genome and its LTR-RTs are allowed to mutate with only DNA substitutions. Three reps with “`-TiTv 10`” and “`-TiTv 1`” allowed us to compare the performance of PrinTEs models of evolution. For comparison, we used SLiM to evolve a single 100 Mb diploid lineage (N=1). SLiM does not implement an iterative stepwise approach and instead models each generation. To reduce runtime, we used SLiM to model ancestral trees with no mutations (MU=0), then projected DNA substitutions onto the mutation-free SLiM genealogies at half-million generationsteps from 0 - 7.5 million generations using a mutation rate of 3×10^-8^ per bp per generation and Ti/Tv’s of 1 or 10. Mutations in SLiM simulated genomes were detected by whole-genome alignments using Minimap2 with flags “`-cx asm20 --cs=long -- secondary=no`”. The PAF alignment files were provided to “`paftools.js`” from the Minimap2 package with the “`stat`” command to calculate p-distance from the reported number of substitutions and alignment length. Similarly, K2P divergence was calculated using the cs tag in the PAF file by classifying substitutions as either transitions (purine to purine or pyrimidine to pyrimidine) or transversions (purine ↔ pyrimidine).

LTR-RT redetection was compared between SimulaTE, ReplicaTE, and PrinTE simulations. SimulaTE requires a chassis genome as input. PrinTE was used to build the chassis using “`--burnin_only --cds_percent 0 --TE_percent 0 --chr_number 1 --size 1Gb`”. The “`define-landscape_random-insertions-freq-range.py`” from SimulaTE was run using “`--N 3 --insert-count 2000 --min-distance 1000 -- min-freq 0.1 --max-freq 0.9`”. The resulting pgd file was provided to “`build-population-genome.py`” from SimulaTE to create the final TE-infused genome. ReplicaTE was used to insert Ty1 and Ty3 elements from *Drosophila melanogaster* chromosome 2 using the GenBank file from the package. PrinTE was run with parameters “`--burnin_only -- cds_percent 0 --TE_num 1000 --chr_number 1 --size 500Mb -s 21 -- threads 200`”. Three replicates were used for each tool. LTR-RT redetection was conducted using `LTR_FINDER_parallel` and `LTR_HARVEST_parallel` with parameters “`-w 2 -C -D 15000 -d 100 -L 7000 -l 60 -itp 20 -M 0.85 -S 3`” and “`-minlenltr 60 - maxlenltr 7000 -mintsd 4 -maxtsd 8 -mindistltr 300 -maxdistltr 20000 - similar 85 -vic 10 -seed 20 -seqids yes -xdrop 6`”, respectively. The resulting LTR-RT candidates were provided to LTR_retriever with “`-motif [TCCA TGCT TACA TACT TGGA TATA TGTA TGCA TGAA AGTT AGCT]`” to identify LTR-RTs.

PrinTE runtime and peak memory were assessed on an HPC server with 256 CPU threads of AMD EPYC 7763 2.45GHz and 1024 GB DIMM DDR4 memory at 3200 MHz. Metrics were collected from the report generated using “`/usr/bin/time -v`”. PrinTE was run using “`-cn 20 -sz 100Mb -P 20`” to simulate a 100Mb genome with 20 chromosomes in which 20% of the genome is genes. “`-itn 6000`” was used to simulate 6000 initial TE insertions. “`-ge 1000000 -st 100000`” simulated 1,000,000 generations of TE evolution with step sizes of 100,000 generations (10 iterations total). Insertion and deletion rates, “`-F 3.0e-11,5e-11`”, were used to insert and delete ∼500 TE insertions per iteration.

### Syntenic Ribbon Plot and LTR-RT Phylogeny

To simulate genomes presented in **Fig. 3A**, 19,621 *Arabidopsis thaliana* cds sequences were inserted into the simulated genome using PrinTE with “`--cds_num 19621`” to guide collinear anchoring and enable syntenic analysis. The burn-in genome consisted of 80% intact LTR-RTs (`-itp 80`). The genome contraction phase was run using “`-ge 5000000 -st 1000000 -F 1e-11,4e-11 -dg -k 3`”. The genome size increase phase was run using identical parameters except “`-F 4e-11,1e-11 –continue`” and the contraction phase outputs as the input. The simulated genomes were processed pairwise using “`jcvi.compara.catalog ortholog -- cscore=.99`” in JCVI (v1.5.4) ^48^. Syntenic anchors were visualized using “`matplotlib.pyplot`” in Matplotlib (v3.10.3) to create the ribbon plot (**Fig. 3A**). The LTR-RT phylogeny (**Fig. 3D,E**) was created using only elements in the *Tekay* family, which were classified as such by TEsorter (v1.4.7) ^49^. *Tekay* LTR-RT headers were tagged with “`_shared`” and “`_uniq`” to facilitate post-PrinTE parsing. Shared LTR-RTs were inserted using PrinTE with parameters “`-itp 80 -cn 1 -sz 10Mb -tmx 10 --TE_lib Tekay_shared.fa -tk 15`”. The outputs (“`burnin.fa`” and “`burnin.bed`”) were duplicated and allowed to continue simulation using either genome expansion parameter ( “`--generation_end 5000000 --step 500000 -F 2e-10,1e-10 --TE_lib Tekay_uniq.fa`”) or genome contraction parameters ( “`--generation_end 5000000 -- step 500000 -F -F 2e-11,1e-11 --TE_lib Tekay_uniq.fa`”). Intact LTR-RTs were extracted using “`extract_intact_LTR.py`” and classified using TEsorter with “`-db rexdb-plant -dp2`”. Reverse transcriptase domains identified from *Tekay* LTR-RTs were aligned using MAFFT (v7.525) ^50^ and provided to VeryFastTree (v4.0.5) ^51^ for phylogenetic analysis. The resulting newick trees were plotted using ggtree (v3.1.5) ^52^ with branches colored by their ancestral relationship.

### *Pucciniomycotina* Genomes and Annotations

The myrtle rust (*Austropuccinia psidii*) genome was downloaded from NCBI with accession ID ERS3953144 ^3^. Genomes of the *Pucciniomycotina* fungus subdivision were obtained from the JGI Fungal Genomics Resource MycoCosm (https://mycocosm.jgi.doe.gov/) (**Table S1**). Rust fungus genomes with contig N50 values higher than 100 kbp were used, for a total of 47 genomes (**Table S2**).

Benchmarking Universal Single-Copy Orthologs (BUSCOs) evaluations were performed on each genome by compleasm (v0.2.7) ^37^ with the odb12 database and the basidiomycota lineage that has 2409 universal single-copy orthologs. For genomes with duplicated BUSCO higher than 5%, the Redundans (v2020.01.28) ^36^ program was used to identify alternative haplotypes. Haplotype alignments with an identity higher than 90% and coverage of more than 80% were removed using “`--identity 0.9 --overlap 80`” parameters. compleasm was again used to evaluate the purged genomes with the same parameter. The resulting duplicated BUSCO were considered biological duplications (**Table S2**). The haplotype genomes (de-duplicated) were used in the following analyses.

Genomes with single-copy BUSCOs higher than 50% were used to construct the phylogeny. Melame1 (BUSCO S:1134,D:1147,F:33,I:0,M:95) and MecolCla_1 (BUSCO S:826,D:1452,F:34,I:0,M:97) were not included in this analysis owing to the high proportion of duplicated BUSCO. The protein sequences of single-copy BUSCOs present in the remaining 45 genomes were extracted and further aligned using MAFFT (v7.525) ^50^ with a focus on local alignments using the “`--maxiterate 1000 --localpair`” parameters. The phylogeny was constructed using the aligned BUSCO and IQ-TREE (v2.4.0) ^38^. Each BUSCO alignment was treated as an individual partition ^53^, and ModelFinder ^54^ was used to identify the best model for each partition using the “`-m MFP`” parameter in IQ-TREE. The Shimodaira–Hasegawa-like approximate likelihood ratio test (SH-aLRT) ^55^ and bootstrap tests were performed 1000 times each with parameters “`-bb 1000 -alrt 1000`” in IQ-TREE. The bootstrap consensus tree was plotted by ggtree (v3.1.5) ^56^ in R.

Transposable elements in each fungal genome were annotated using EDTA (v2.2.2) ^57^ following the instructions of Benson et al. (2024) ^58^. The “`--force 1`” parameter was used to overcome analysis in genomes with missing types of TEs (*i*.*e*., LTR-RTs). The “`--anno 1`” parameter was used to obtain whole-genome TE annotations. The “`--cds`” parameter was used to provide coding sequences to EDTA to improve TE annotations. Primary coding sequences of each genome were obtained from JGI MycoCosm and Tobias et al. (2021) ^3^. Solo and intact LTR-RTs were identified from whole-genome TE annotations using LTR_retriever (v3.0.1) ^30^. The solo:intact ratio was calculated on the whole-genome level, which is the number of whole-genome solo LTRs divided by the number of whole-genome intact LTR-RTs. “inf” values were given to genomes without intact LTR-RTs. The divergence of intact LTR-RTs was estimated by LTR_retriever (v3.0.1) ^30^ using the one-parameter Jukes-Cantor 69 model ^59^.

### Simulating Digital Replicas of Rust Fungus Genomes

Three fungal genomes were chosen for simulations, including *Puccinia coronata* f. sp. *avenae* (PuccoNC29_1), *Cronartium ribicola* (Crori1), and *A. psidii* (myrtle rust). TE libraries used for TE removal and simulating the fungal genomes were obtained via *de novo* TE annotation using EDTA (as described above). The Genome Approximation Mode of TEgenomeSimulator (v1.0.0) was used for simulations. The resulting digital replicas of the rust fungus genomes were further used to simulate LTR dynamics using PrinTE (see sections below).

### Forward Simulation of Rust Fungus Genomes

#### Simulating LTR-RT Dynamics in Rust Fungus Genomes

##### Cronartium ribicola

We hypothesized that this genome is relatively stable, with high rates of LTR-RT insertions and deletions. We aimed to evolve the genome with a relatively stable genome size. The digital replica of the *C. ribicola* genome was generated by TEGenomeSimulator with a size of 232.6 Mb, which was used as input for PrinTE using the ‘`--bed`’ and ‘`--fasta`’ parameters. Each experiment was simulated with 20,000 generations per step and a total of 200,000 generations (`-ge 200000 -st 20000`). A series of fixed insertion and deletion rates was used in different experiments, including the range of insertion rates from 1.0×10^-9^ to 1.9×10^-8^ per bp per generation and the range of deletion rates from 1.4×10^-9^ to 4.0×10^-8^ per bp per generation specified by the ‘`--fix`’ parameter. Additionally, DNA substitution rates of 1.0×10^-8^, 2.0×10^-9^, and 5.0×10^-9^ per bp per generation, k values of 0, 3, 6, and 10, and solo rates of 60, 70, 75, 80, and 95 were used in the parameter sweep analysis. The simulated genomes were summarized by PrinTE and LTR_retriever and compared with the real *C. ribicola* genome for the determination of the best parameters.

##### *Puccinia coronata* f. sp. *Avenae*

We hypothesized that this genome is relatively stable with low rates of LTR-RT insertions and deletions. We aimed to evolve the genome with a relatively stable genome size. The digital replica of the *P. coronata* f. sp. *avenae* genome was generated by TEGenomeSimulator with a size of 112.7 Mb, which was used as input for PrinTE using the ‘`--bed`’ and ‘`--fasta`’ parameters. Each experiment was simulated with 1 million generations per step and a total of 10 million generations (`-ge 10000000 -st 1000000`). A series of fixed insertion and deletion rates was used in different experiments, including the range of insertion rates from 8.0×10^-12^ to 5.0×10^-11^ per bp per generation and the range of deletion rates from 9.6×10^-12^ to 9.0×10^-11^ per generation specified by the ‘`--fix`’ parameter. Additionally, mutation rates of 2.0×10^-9^ and 5.0×10^-9^ per bp per generation, k values of 0, 3, and 6, and solo rates of 70 and 95 were used in the parameter sweep analysis. The simulated genomes were summarized by PrinTE and compared with the real *P. coronata* f. sp. *avenae* genome for the determination of the best parameters.

##### Austropuccinia psidii

We hypothesized that this was once an expanding genome but now a fast downsizing genome with a low rate of LTR-RT insertion and a high rate of LTR-RT deletion. Owing to the relative phylogenetic closeness to *A. psidii*, we use the genome size and TE composition of *P. coronata* f. sp. *avenae* genome as an approximation of the ancestral state of *A. psidii* before genome expansion. We used PrinTE to simulate such an ancestral genome with parameters ‘`--cds_percent 20 --TE_num 6000 --chr_number 20 --size 113Mb --TE_mut_k 10 --TE_mut_Mmax 20`’. With a fixed deletion rate of 5.0×10^-13^ per bp per generation, we explored insertion rates from 2.0×10^-12^ to 9.8×10^-11^ per bp per generation and identified that the insertion rate of 1.4×10^-11^ per bp per generation could yield LTR divergence similar to the expansion phase of *A. psidii* (**Fig. S13A**,**B**). We thus used PrinTE to simulate a fast-expanding phase with parameters “`-k 0 --fix 1.4e-11,5e-13 --mutation_rate 2e-9 --solo_rate 95`” until it reached 1,178 Mb. For the contraction phase, we slowed down the insertion rate to 1.4×10^-13^ per bp per generation and explored deletion rates from 3.0×10^-13^ to 5.0×10^-11^ per bp per generation. We identified that the deletion rate of 3.0×10^-11^ per bp per generation could yield LTR divergence and genome size similar to those of the *A. psidii* genome (**Fig. S13A**,**B**). Finally, we used the outputs of the last iteration as inputs for PrinTE to simulate a fast-contracting phase with parameters “`-k 0 --fix 1.4e-13,3e-11 -- mutation_rate 2e-9 --solo_rate 95`” until it reached 1000 Gb, approximately the size of the *A. psidii* genome.

## Data Availability

All codes that produced the findings of the study, including all main and supplemental figures, are available at https://github.com/cwb14/PrinTE.git, under the GPL-3.0 license. The Kmer2LTR code is available at https://github.com/cwb14/Kmer2LTR.git under the GPL-3.0 license.

## Acknowledgments

We want to thank Drs. Jason Slot, James Pease, and H. Lisle Gibbs for helpful discussions of this project. We are also grateful to Drs. Chen Wu, Julie Blommaert, Olivia Angelin-Bonnet, and David Chagné for reviewing this manuscript and providing insightful comments.

## Contributions

S.O. conceived the project and supervised the work. C.W.B developed PrinTE and Kmer2LTR. C.W.B, T.-H.C., and S.O. performed the analysis. C.W.B, T.-H.C., C.H.D, S.J.T., and S.O. wrote the manuscript text. All authors reviewed and approved the manuscript.

## Ethics declarations

### Ethics approval and consent to participate

Not applicable.

