## Supplemental Figures for "PrinTE: A Forward Simulation Framework for Studying the Role of Transposable Elements in Genome Expansion and Contraction"

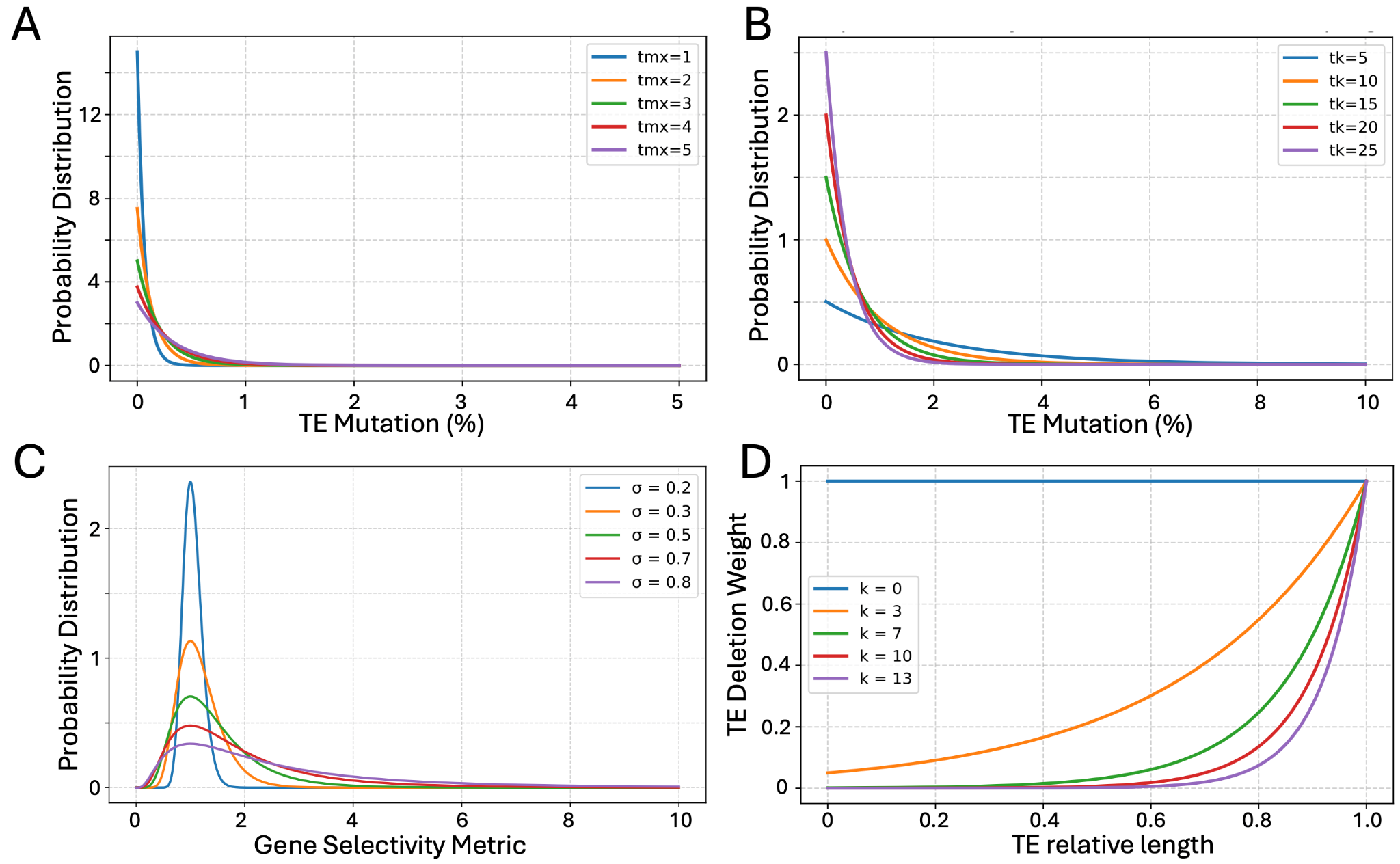

**Fig. S1.** PrinTE evolutionary modelling based on customizable probability distributions. The distribution of mutation probabilities for TEs inserted into the burn-in (generation zero) genome is modulated by an exponential decay function, where maximum mutation of TEs (‘–-tmx’; **A**) and TE mutation bias (‘--tk’; **B**) are used to customize the distribution. (**C**) Gene selectivity is scored using a log-normal distribution where σ controls the distribution of selective scores. The higher the Gene Selectivity Metric value, the higher the selection coefficient of TE insertions into genes. A narrower density distribution indicates that most gene mutations by TEs are near neutral or only slightly deleterious. (**D**) The element length determines TE’s probability of deletion, and selectivity is determined by the -k parameter. A higher k will impose a higher weight on deleting long TEs. k=0 means that all TEs have an equal probability of being deleted.

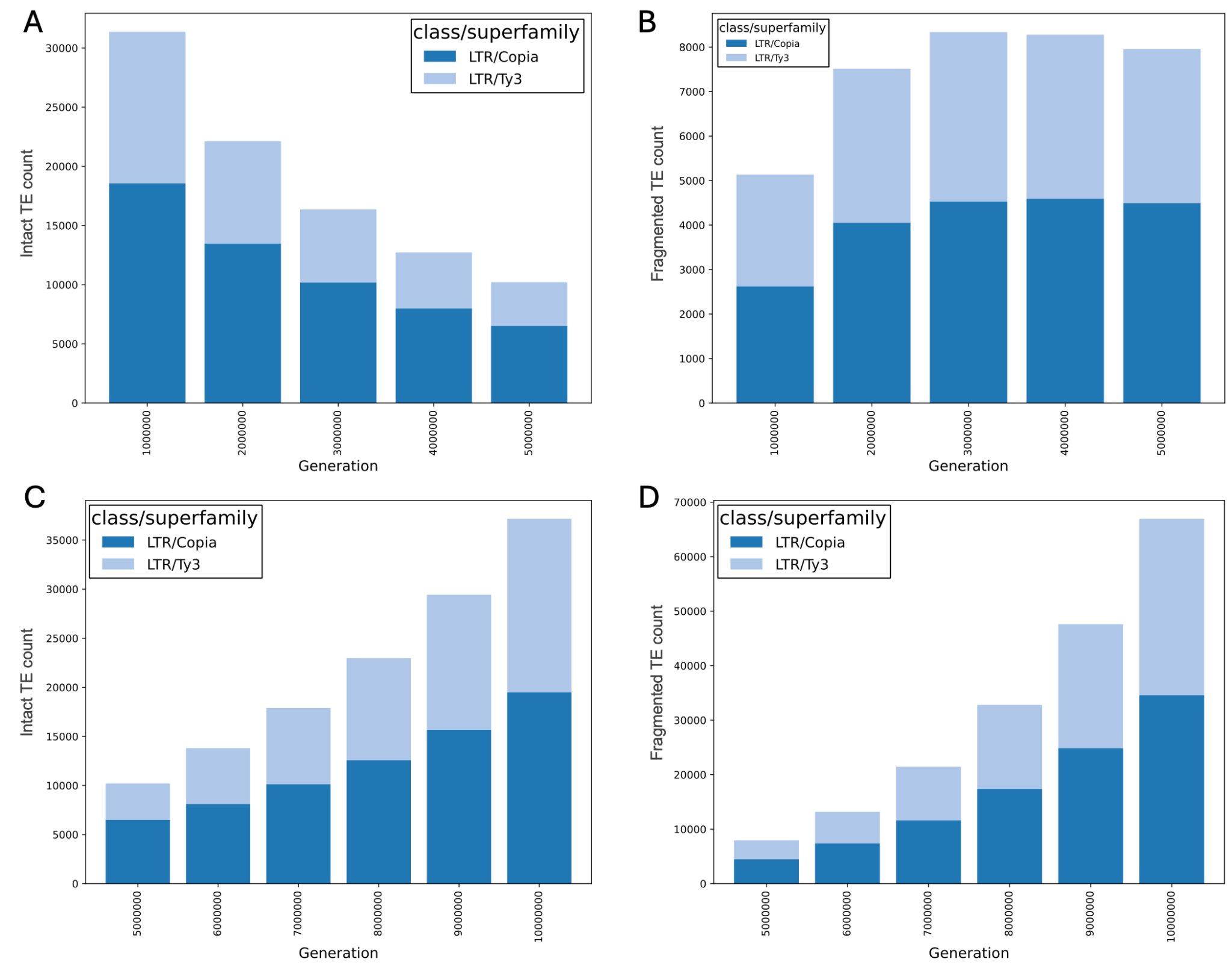

**Fig. S2.** TE counts across generations in PrinTE simulations. Intact (**A**) and fragmented (**B**) TE counts in a simulated genome experiencing a genome size decrease. Intact (**C**) and fragmented (**D**) TE counts in a simulated genome experiencing a genome size increase. The data are the same as those used in **Fig. 3**. (**A**) and (**B**) correspond to the first five million generations, and (C) and (D) correspond to the second five million generations.

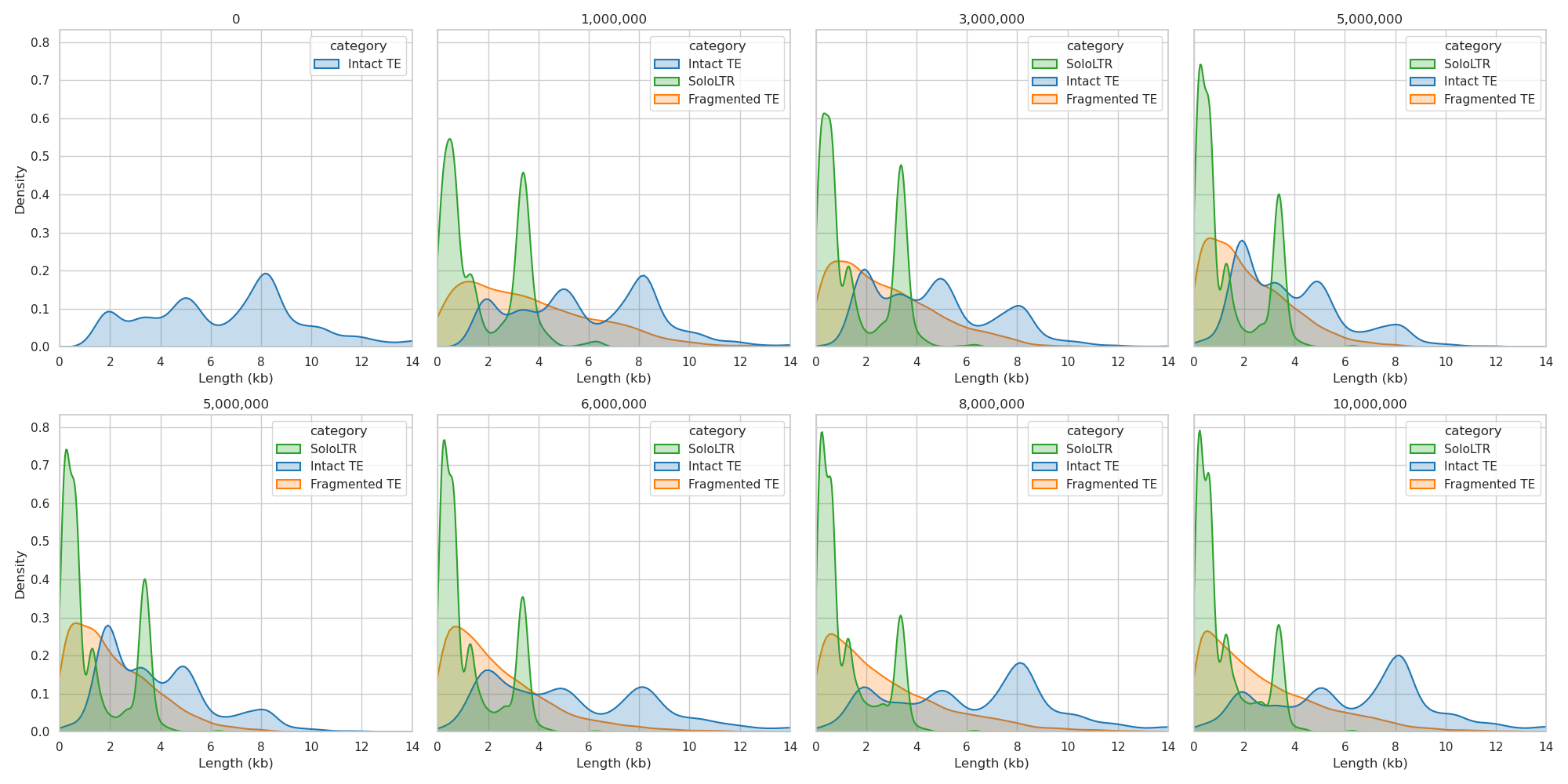

**Fig. S3.** The density of PrinTE feature lengths for intact and fragmented LTR-RTs and solo LTRs. The top row shows a period of genome contraction, characterized by progressive buildup of solo LTRs and fragmented LTRs (zero to 5 million generations are shown). The bottom shows feature density during a subsequent period of genome expansion. The abundances of solo LTRs and fragmented LTRs are maintained, but the proportion of longer and intact LTR-RTs increases through time.

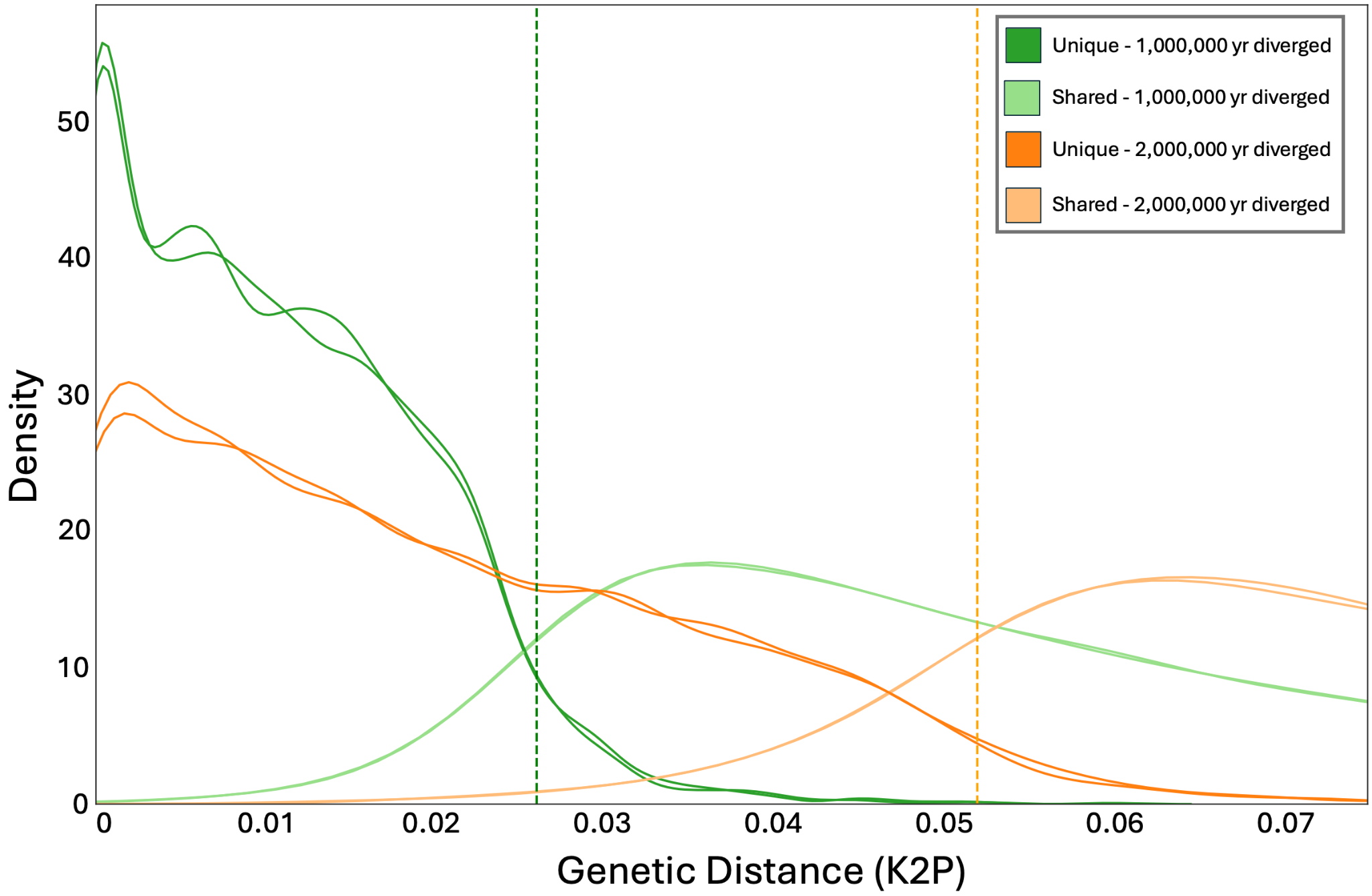

**Fig. S4.** PrinTE simulated LTR-RT divergence between two lineages after one million (green) and two million (orange) generations after reproductive isolation. Shared LTR-RTs are retained in both lineages and derived from existing TEs of the most recent common ancestor (MRCA). Unique LTR-RTs are only present in one of the lineages and were transposed after lineage divergence. The vertical dashed line shows the expected genetic distance of the MRCA at each time point.

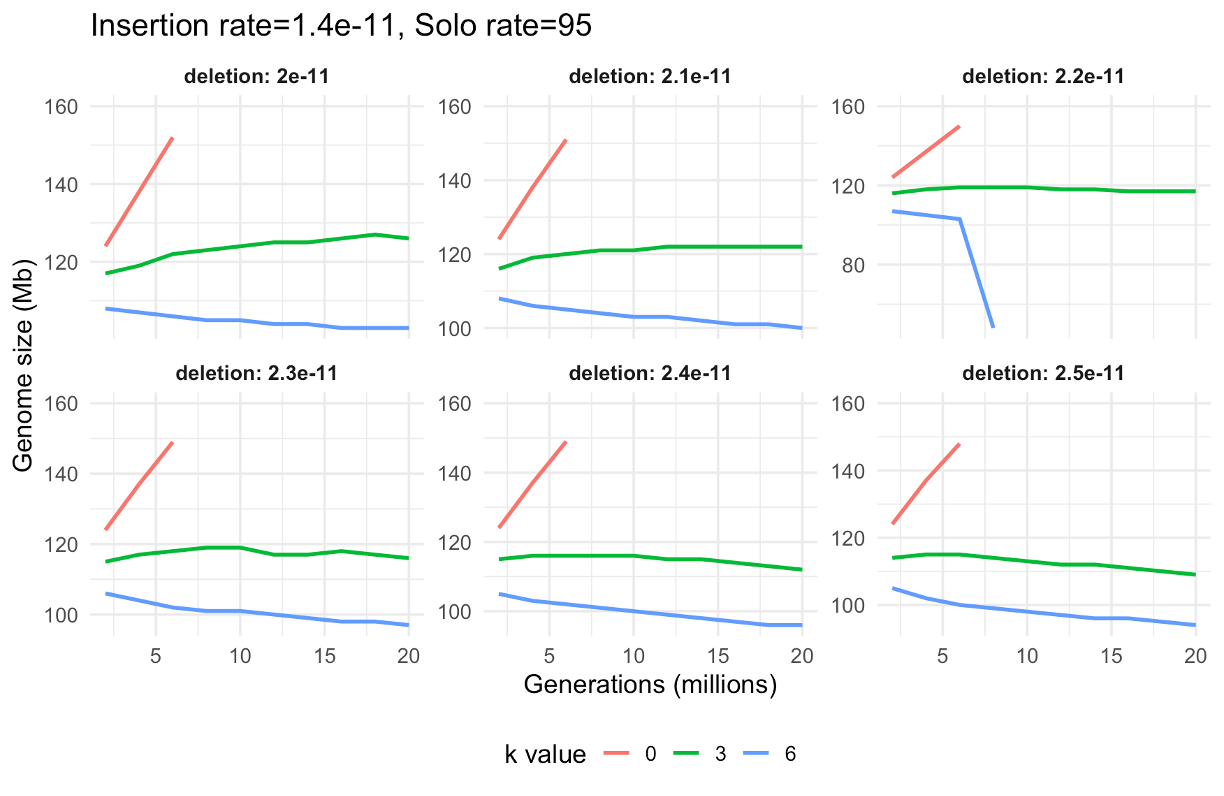

**Fig. S5.** Variation of genome size simulated using PrinTE under fixed insertion rate (1.4×10^-11^ /bp/generation), varying deletion rates (2×10^-11^ – 2.5×10^-11^ /bp/generation), varying deletion length bias (k=0, 3, or 6), and fixed rates of deletion events creating solo LTRs (95%).

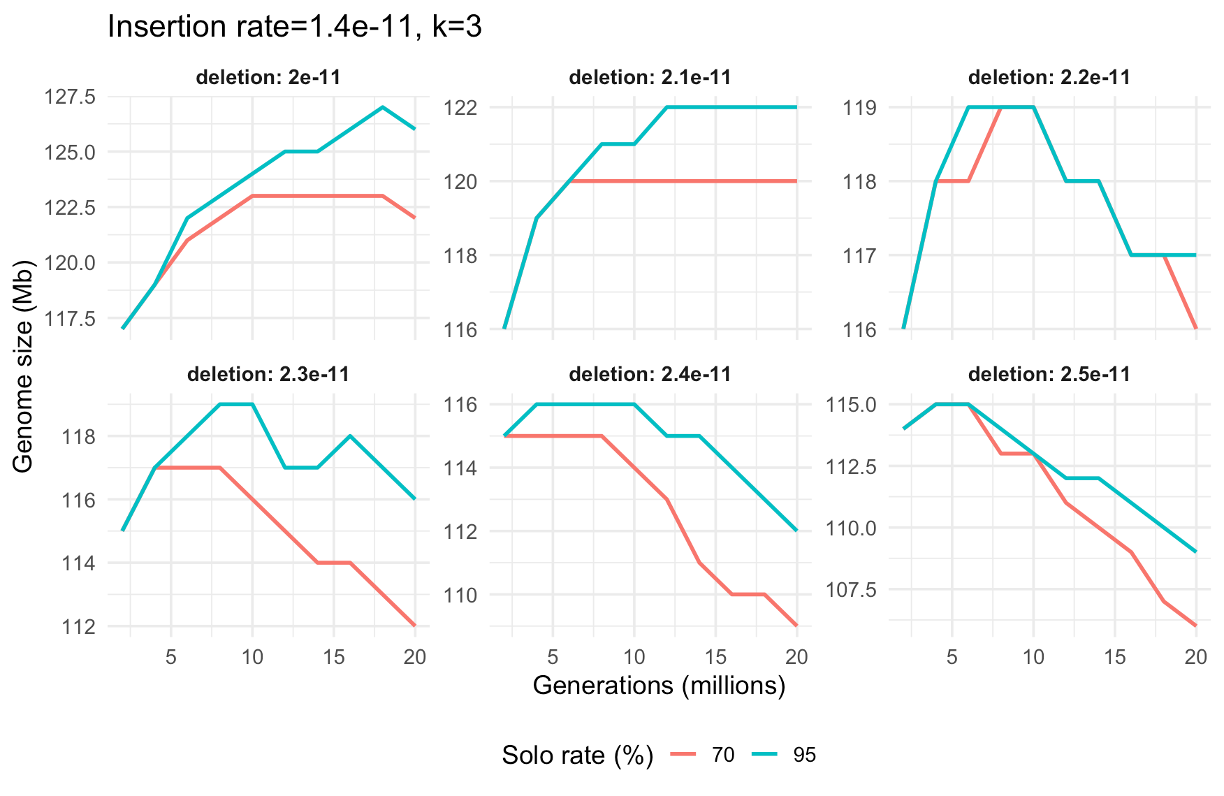

**Fig. S6.** Variation of genome size simulated using PrinTE under fixed insertion rate (1.4×10^-11^/bp/generation), varying deletion rates (2×10^-11^ – 2.5×10^-11^ /bp/generation), fixed deletion length bias (k=3), and varying rates of deletion events creating solo LTRs (70% and 95%).

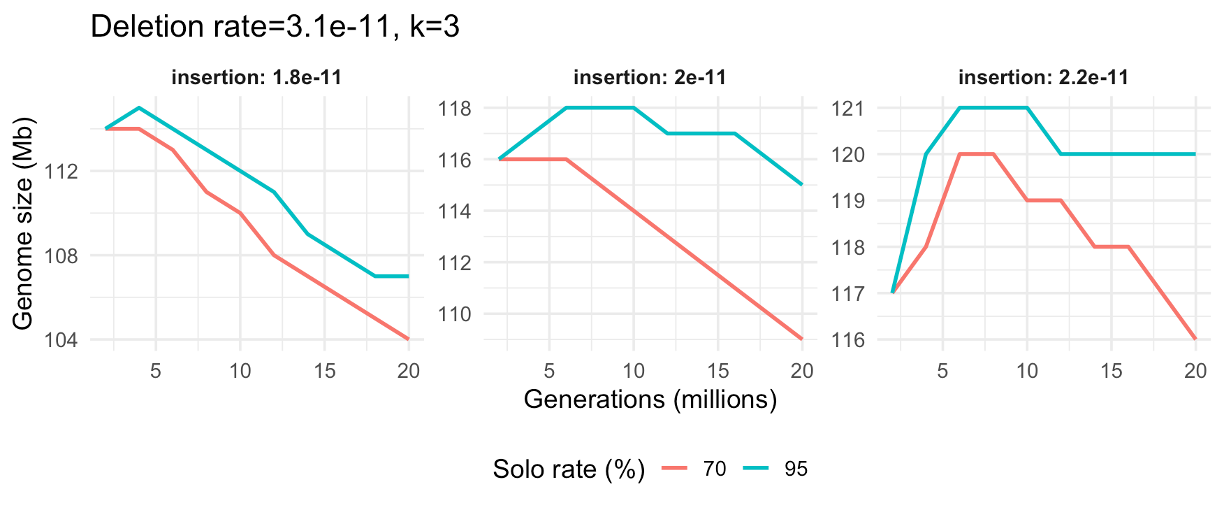

**Fig. S7.** Variation of genome size simulated using PrinTE under varying insertion rate (1.8×10^-11^- 2.2×10^-11^ /bp/generation), fixed deletion rates (2×10^-11^ – 2.5×10^-11^ /bp/generation), fixed deletion length bias (k=3), and varying rates of deletion events creating solo LTRs (70% and 95%).

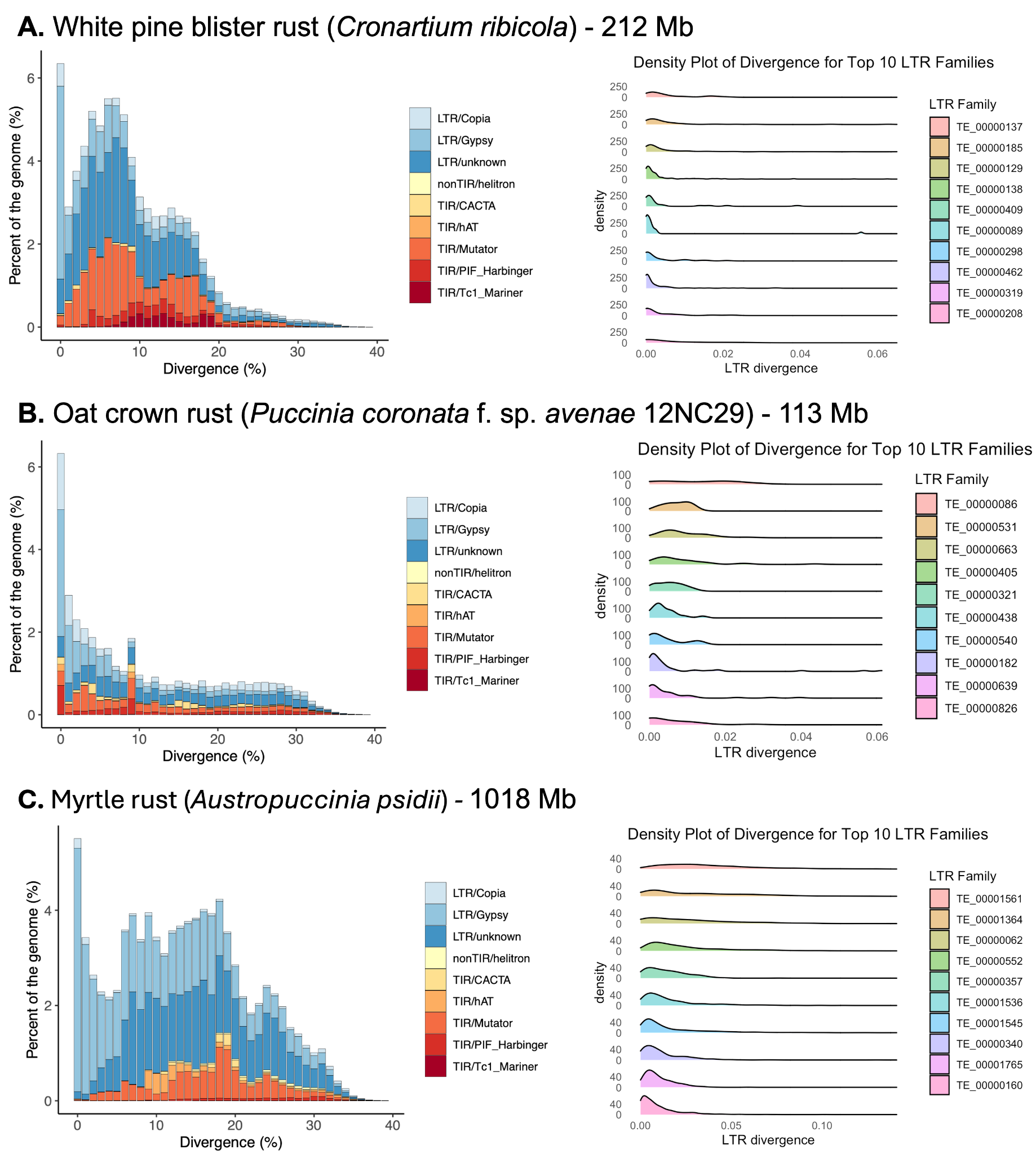

**Fig. S8.** Transposable element divergence in *Cronartium ribicola* (**A.** white pine blister rust), *Puccinia coronata* f. sp. *avenae* (**B.** oat crown rust), and *Austropuccinia psidii* (**C.** myrtle rust) genomes. The divergence distribution of the top 10 most abundant intact LTR families in each genome is shown, and they are not shared between genomes.

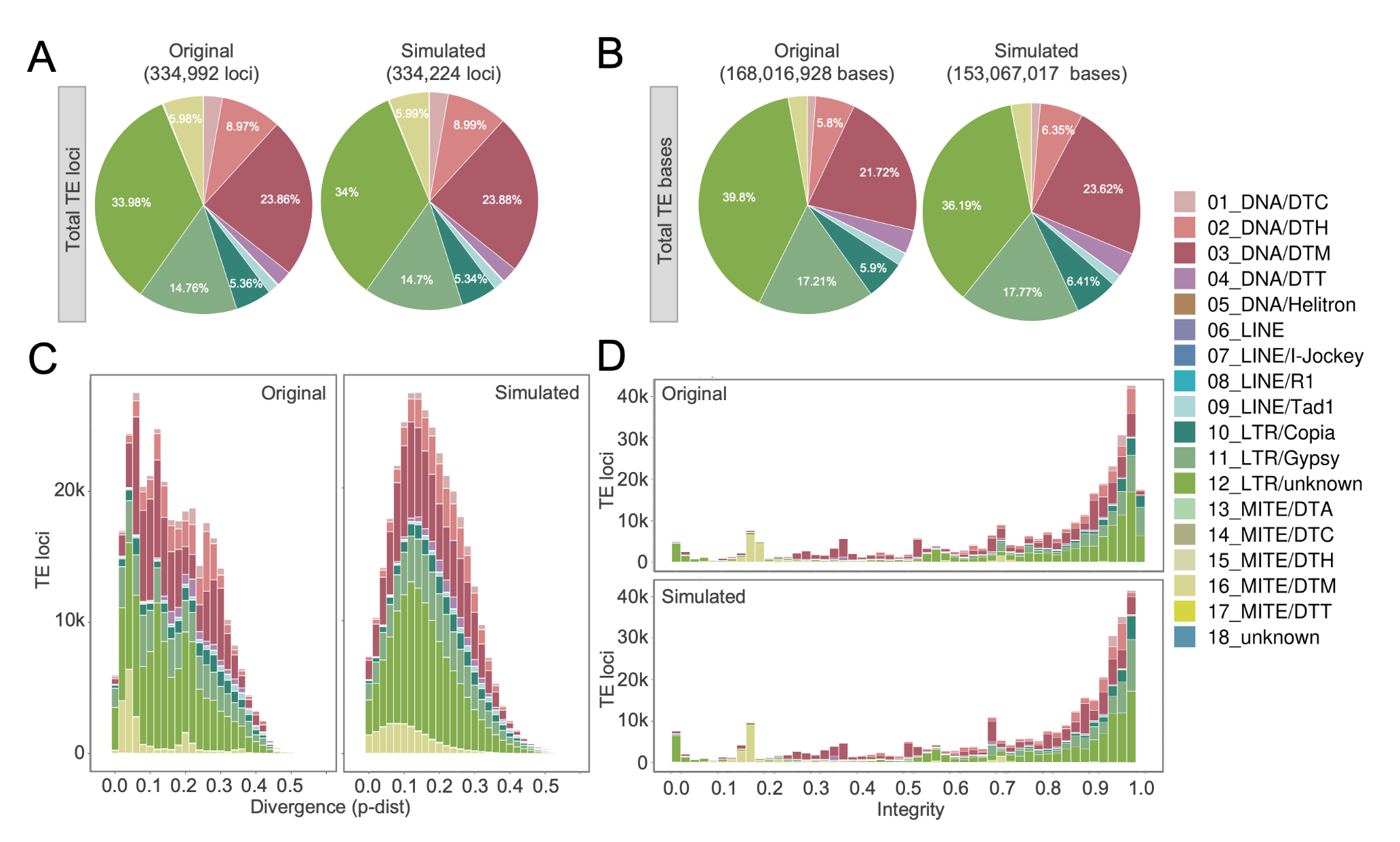

**Fig. S9.** **TE Composition Approximation for the *Cronartium ribicola* genome.** (**A**) Comparison of total TE loci between the original and simulated genomes. (**B**) Comparison of total TE bases between original and simulated genomes. (**C**) TE sequence diversity distribution colored by TE superfamily. (**D**) The distribution of TE sequence integrity, which is the length ratio relative to the consensus sequence, is colored by TE superfamily as in (**C**).

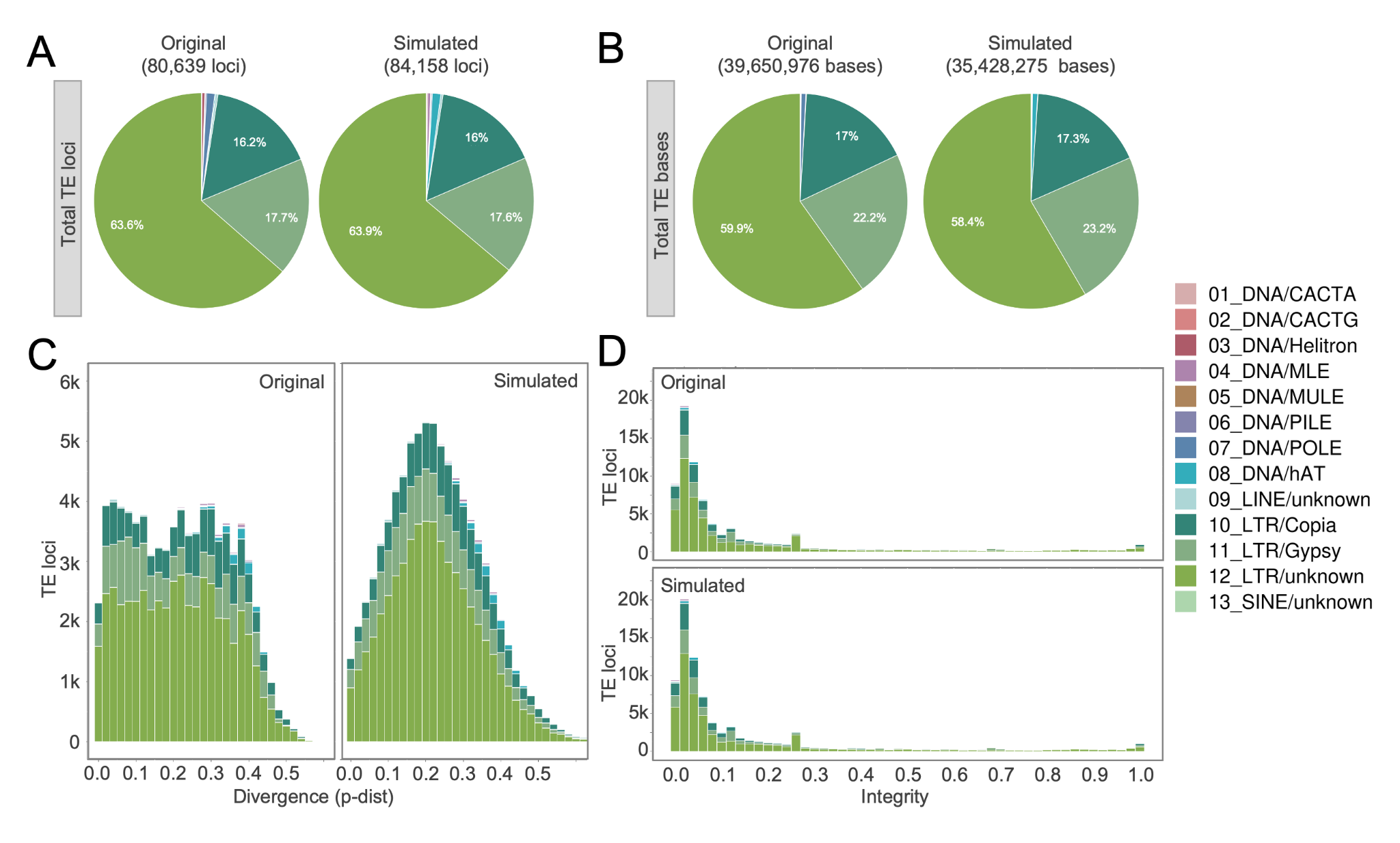

**Fig. S10.** **TE Composition Approximation for the *Puccinia coronata* f. sp. *avenae* genome.** (**A**) Comparison of total TE loci between the original and simulated genomes. (**B**) Comparison of total TE bases between original and simulated genomes. (**C**) TE sequence diversity distribution colored by TE superfamily. (**D**) The distribution of TE sequence integrity, which is the length ratio relative to the consensus sequence, is colored by TE superfamily as in (**C**).

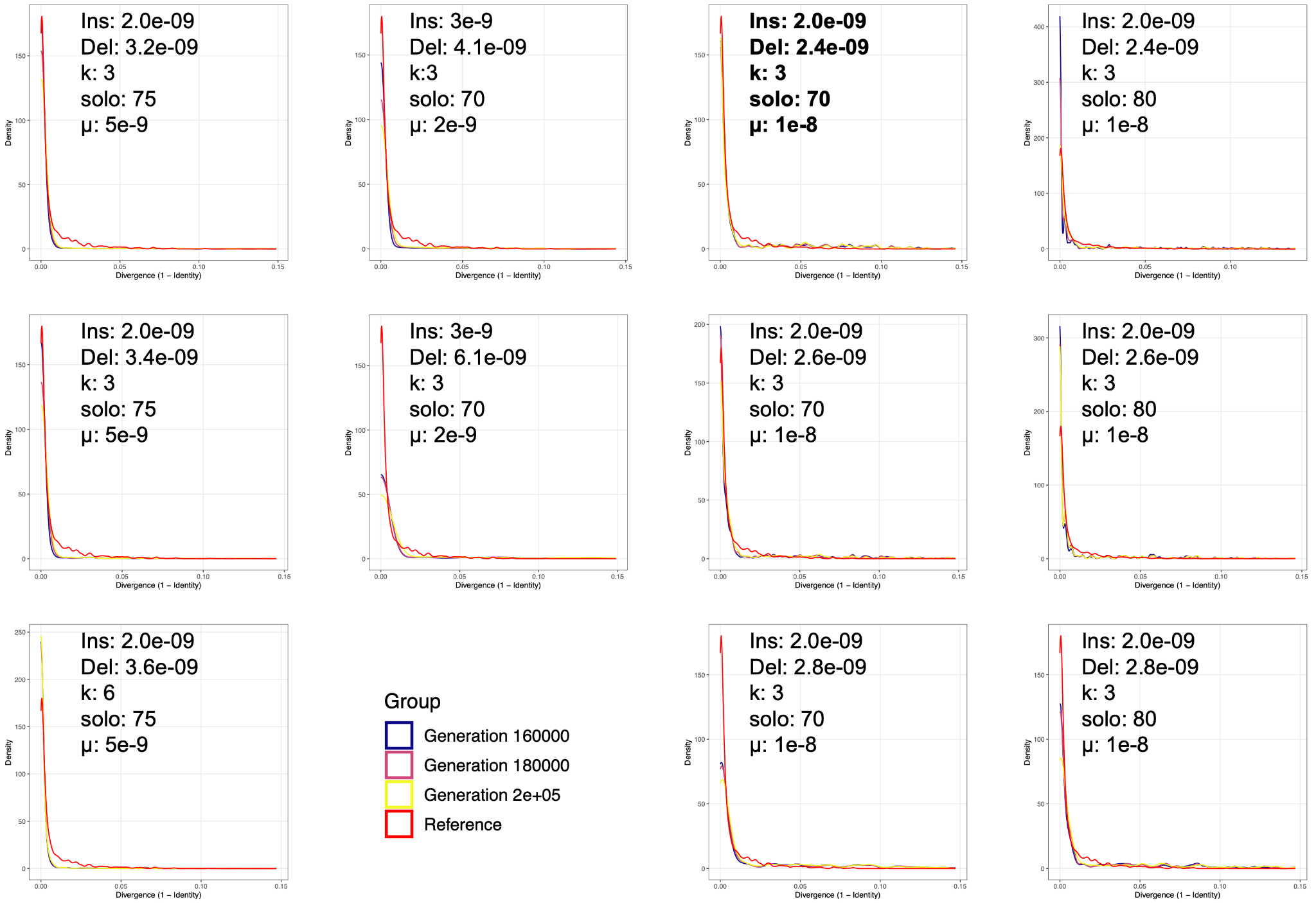

**Fig. S11.** Density distributions of LTR-RT divergences using the Kimura two-parameter (K2P) model that compare the *Cronartium ribicola* genome (red lines) to PrinTE simulations. Blue, purple, and yellow lines show LTR-RT divergences sampled from three forward simulation time points. The parameters in bold were selected as they generated the distribution most closely mimicking that of *C. ribicola.*

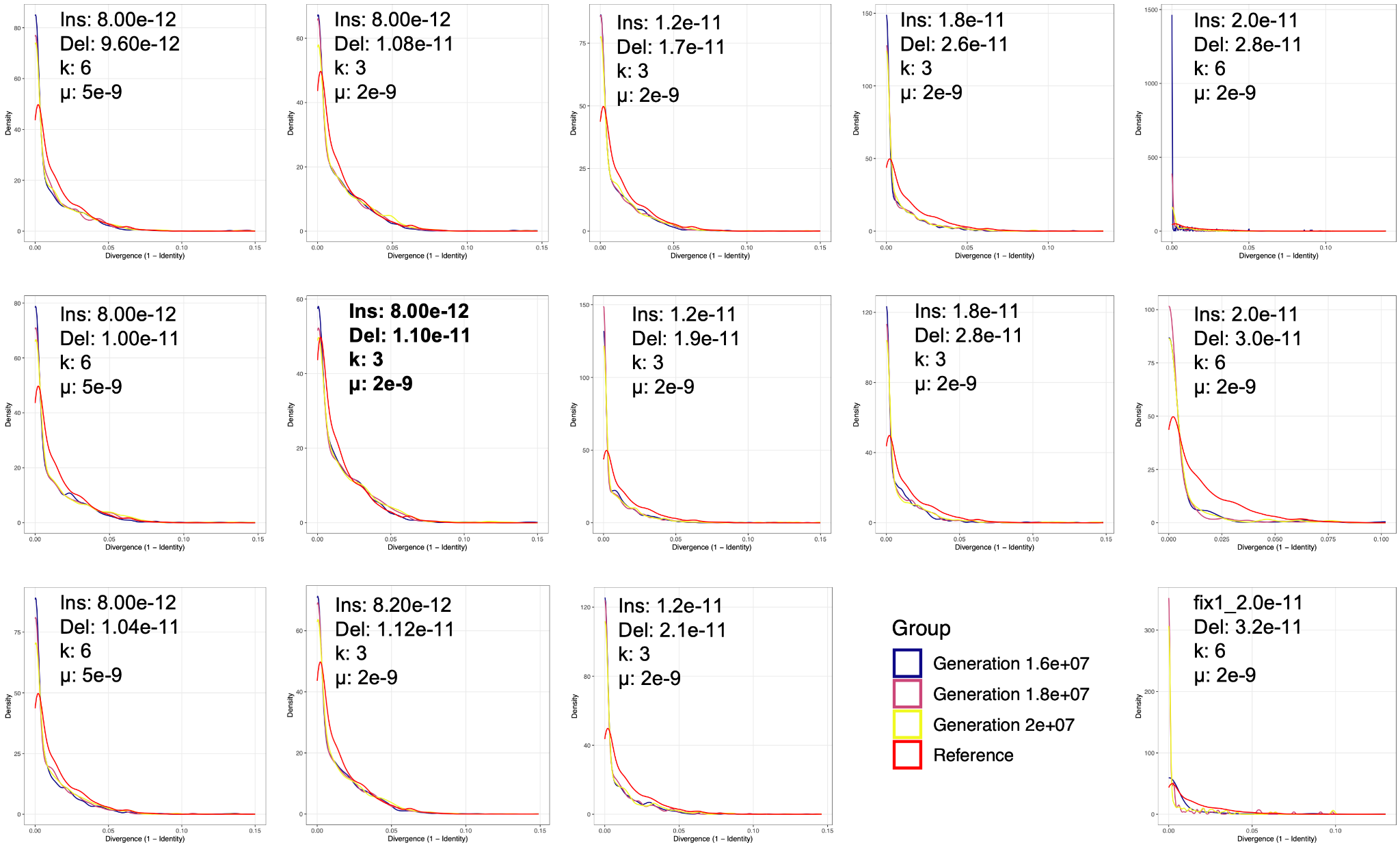

**Fig. S12.** Density distributions of LTR-RT divergences using the Kimura two-parameter (K2P) model that compare the *Puccinia coronata* f. sp. *avenae* genome (red lines) to PrinTE simulations. All simulations shown used the solo rate of 70% (--solo_rate 70). Blue, purple, and yellow lines show LTR-RT divergences sampled from three forward simulation time points. The parameters in bold were selected as they generated the distribution most closely mimicking that of *P. coronata.*

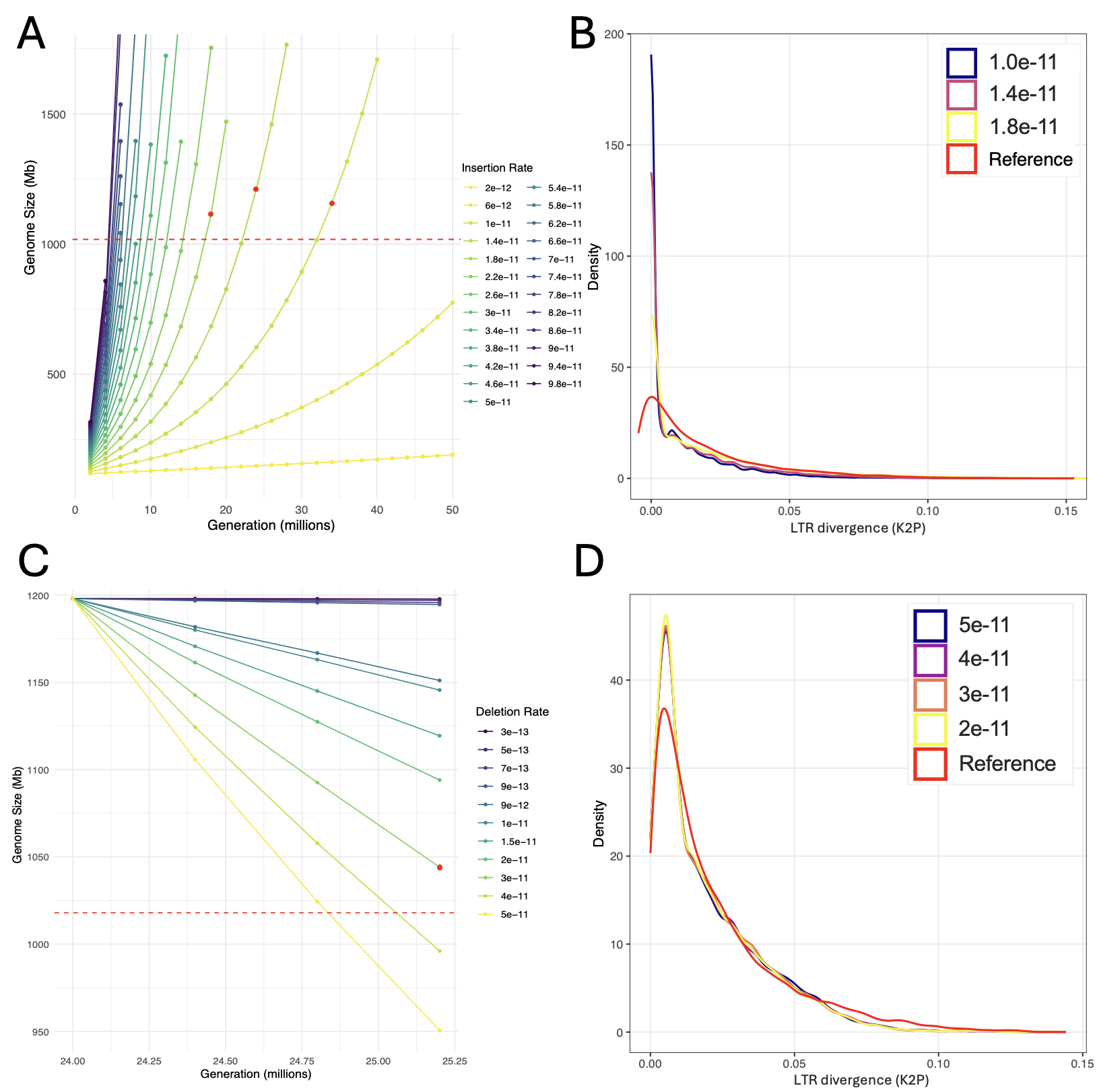

**Fig. S13**. The method to compare *Austropuccinia psidii* (myrtle rust) simulations. The *A. psidii* simulation occurs in two phases: a long phase of genome expansion (**A-B**) and a short phase of genome contraction (**C-D**). (**A**) Genome size change using varying insertion rates to simulate the genome expansion phase. A fixed deletion rate of 5×10^-13^ per bp per generation was used in these simulations. The dashed red line shows the size of the *A. psidii* genome. The three simulations highlighted with red dots are candidates and plotted in **B** to further evaluate. (**B**) Density distributions of LTR-RT divergences using the Kimura two-parameter (K2P) model that compare the *A. psidii* genome (red lines) to PrinTE simulations during the expansion phase. Because these simulations aim to replicate the expansion phase, the reference line was manually shifted left so that the density peak intersects the y-axis. (**C**) The change in genome size using varying deletion rates. A fixed insertion rate of 1.4×10^-11^ per bp per generation was used in these simulations. The dashed red line shows the size of the *A. psidii* genome. The simulation highlighted with a red dot is determined to be the best fit to the real data. (**D**) The density distribution of LTR-RT divergences estimated using K2P for candidate simulations.
